# The Gut Microbiome Controls Liver Tumors via the Vagus Nerve

**DOI:** 10.1101/2024.01.23.576951

**Authors:** Kylynda C. Bauer, Rajiv Trehan, Benjamin Ruf, Yuta Myojin, Mohamed-Reda Benmebarek, Chi Ma, Matthias Seifert, Amran Nur, Jonathan Qi, Patrick Huang, Marlaine Soliman, Benjamin L. Green, Simon Wabitsch, Danielle A. Springer, Francisco J. Rodriguez-Matos, Shadin Ghabra, Stephanie N. Gregory, Jennifer Matta, Brian Dawson, Jihye Golino, Changqing Xie, Amiran Dzutsev, Giorgio Trinchieri, Firouzeh Korangy, Tim F. Greten

## Abstract

Liver cancer ranks amongst the deadliest cancers. Nerves have emerged as an understudied regulator of tumor progression. The parasympathetic vagus nerve influences systemic immunity via acetylcholine (ACh). Whether cholinergic neuroimmune interactions influence hepatocellular carcinoma (HCC) remains uncertain. Liver denervation via hepatic vagotomy (HV) significantly reduced liver tumor burden, while pharmacological enhancement of parasympathetic tone promoted tumor growth. Cholinergic disruption in Rag1KO mice revealed that cholinergic regulation requires adaptive immunity. Further scRNA-seq and in vitro studies indicated that vagal ACh dampens CD8+ T cell activity via muscarinic ACh receptor (AChR) CHRM3. Depletion of CD8+ T cells abrogated HV outcomes and selective deletion of *Chrm3* on CD8^+^ T cells inhibited liver tumor growth. Beyond tumor-specific outcomes, vagotomy improved cancer-associated fatigue and anxiety-like behavior. As microbiota transplantation from HCC donors was sufficient to impair behavior, we investigated putative microbiota-neuroimmune crosstalk. Tumor, rather than vagotomy, robustly altered fecal bacterial composition, increasing Desulfovibrionales and Clostridial taxa. Strikingly, in tumor-free mice, vagotomy permitted HCC-associated microbiota to activate hepatic CD8+ T cells. These findings reveal that gut bacteria influence behavior and liver anti-tumor immunity via a dynamic and pharmaceutically targetable, vagus-liver axis.

## Introduction

Primary liver cancer remains a leading cause of global cancer-related death (*1*, *2*). Furthermore, the liver is one of the most common organs for tumor metastasis (*3*). Hepatic immunosuppression contributes to poor cancer outcomes and exacerbates mortality across different types of cancer (*4*). The liver is a markedly immune tolerant organ (*5*). Continually exposed to gut-derived antigens, the liver actively dampens inflammation (*5*, *6*). This environment facilitates rapid tumor growth and impairs immunotherapy efficacy (*7–9*).

Beyond immune infiltration, nerves within the periphery innervate and shape the tumor microenvironment (*10*, *11*). Direct nerve-tumor signals (*12–14*) and indirect neuroimmune circuits (*15*) likely influence tumor burden. The emerging concept of cancer neuroscience captures these dynamic, bidirectional, and largely unstudied interactions (*16*). Here, we assess how bidirectional vagal activity (vagus ➔ liver, liver ➔ brain) regulates liver tumor outcomes.

The vagus nerve (cranial nerve X) represents the main component of the parasympathetic nervous system, modulating visceral organs via ACh activity (*17*, *18*). Cholinergic vagal activity influences breast (*19*), colorectal (*20*), gastric (*21*), pancreatic (*22*), and prostate tumor progression (*23*). Recently, vagal innervation has been implicated in the prognosis and pathology of liver cancers (*24–26*). Cholinergic nerve density correlated with poor outcomes in HCC patients (*26*) and a retrospective study of nearly 50,000 patients treated for peptic ulcers reported a decreased risk of liver and biliary cancers after truncal vagotomy (abdominal denervation) compared to suture procedures (non-denervation) (*24*). In murine HCC models, non-vagal cholinergic synthesis regulated CD4+ T cell immunosurveillance and exhaustion; genetic ablation of choline acetyltransferase (*Chat*) in CD4+ T cells exacerbated liver tumor burden (*25*). In contrast, cholinergic vagal activity reduced pro-inflammatory cytokine release by macrophages (sepsis models) (*27*) and tumor-associated macrophages (pancreatic cancer models) (*22*). This anti-inflammatory modulation likely shapes the tumor microenvironment (*27*). Whether, and to what extent, this cholinergic anti-inflammatory arc influences hepatic anti-tumor immunity remains uncertain.

We report that surgical hepatic branch vagotomy in mice significantly reduced liver tumor burden. Vagotomized livers exhibited broad inflammatory responses, supporting a cholinergic anti-inflammatory arc (*27*, *28*). Transcriptional and trajectory analyses revealed distinct Ach regulation across CD8+ T cell subsets. We establish that ACh signaling affects CD8+ T cell effector function. Indeed, cholinergic activity impaired CD8+ T cell inflammation, cytotoxicity, and tumor-mediated killing, while genetic ablation of AChR *Chrm3* on CD8+ T cells reduced HCC tumor growth. Compared to sham surgical controls, HV mice also exhibited reduced cancer fatigue and anxiety-like responses, behaviors linked to microbial dysbiosis. HCC microbiota transplantation revealed that gut microbes regulated hepatic anti-tumor immunity via vagal nerves. This gut-brain arc identifies potential targets for neuroimmune-directed therapy and liver cancer treatment.

## Results

### Hepatic vagotomy reduces hepatic tumor burden in mice

To address vagal influence in liver cancer, mice underwent a surgical hepatic vagotomy (HV) or sham procedure (SV) as described (*29*). Severance of the common hepatic vagal branch was confirmed visually and via H&E histology (Fig. 1A). No changes to body and liver weights, as well as liver histology, were observed following hepatic vagotomy (fig. s1A-C).

**Figure 1.**
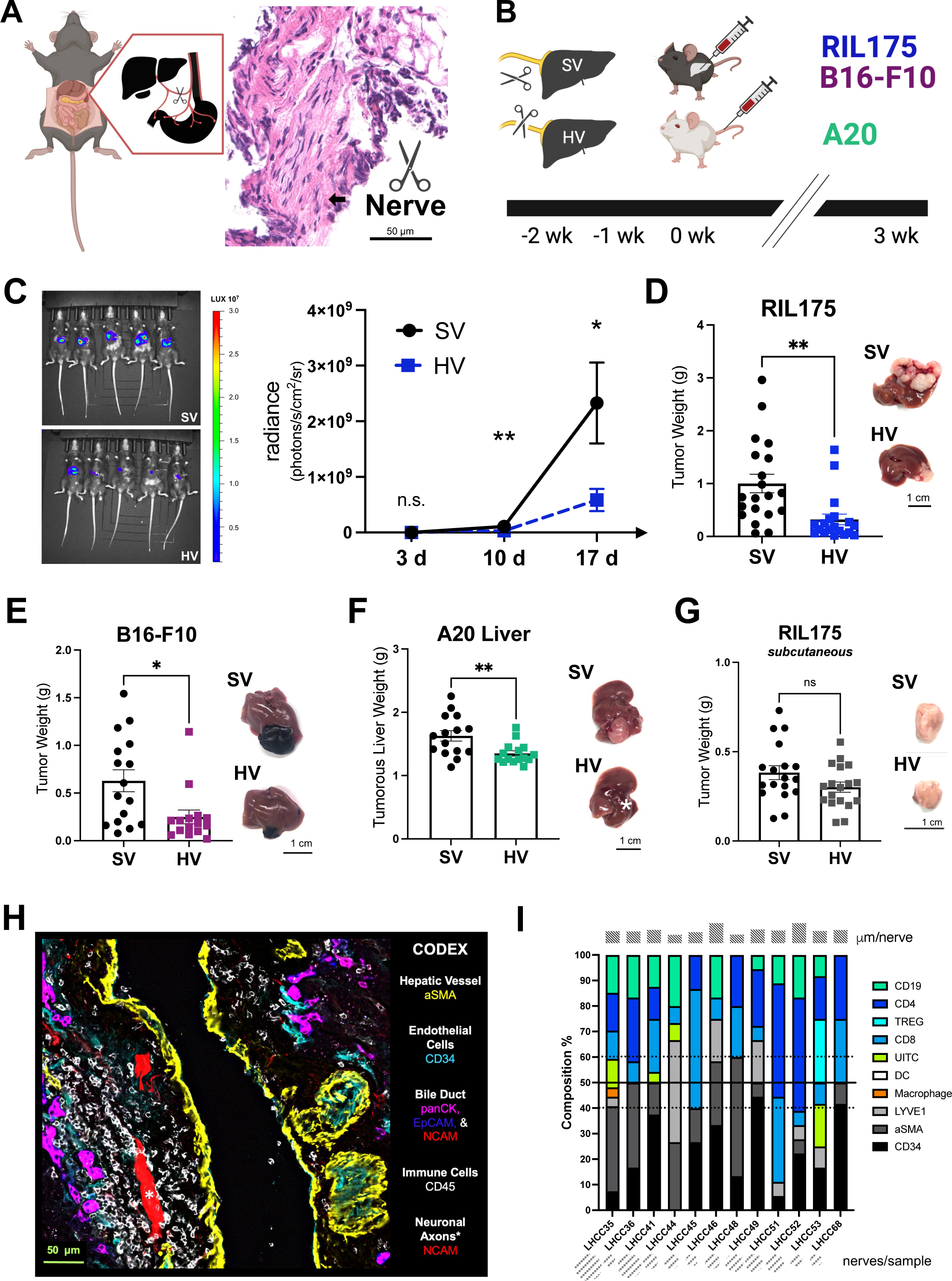
Specific hepatic vagotomy reduces tumor burden in liver cancer. **(A)** Specific hepatic branch vagotomy and innervation diagram with H&E histology of nerve snip. **(B)** Tumor cancer models: C57BL/6 mice underwent intrahepatic injection of RIL175 (hepatocellular carcinoma) or B16-F10 (melanoma) tumor cells following a sham surgical procedure (SV mice) or hepatic vagotomy (HV mice). Alternatively, A20 (lymphoma) cells were injected via tail vein in BALB/c mice. Text color represents tumor models. Unless otherwise noted, tumor burden and hepatic immune composition was assessed 21 d following tumor initiation. (**C**) Representative in vivo imaging (IVIS® Spectrum) from luciferase-expressing RIL175 tumors 10 d following intrahepatic injection. Luminescence across tumor growth (*n* = 10 SV, 9 HV). (**D**) Tumor weight and representative images of RIL175 model. Data includes mice presented in **C**, *n* = 20 SV and 19 HV. (**E**) Tumor weight and representative images of B16-F10 model (*n* = 16 SV, 15 HV). (**F**) Metastatic tumor (METs) diameter and representative images of A20 model (*n* = 15 SV, 15 HV). White star indicates a tumor nodule. (**G**) Subcutaneous RIL175 flank model with tumor weights and representative images (*n* = 18 SV, 18 HV). Data in **D**-**G** represent mean ± SEM of two independent experiments. (**H)** Co-detection by indexing (CODEX) multiplexed immunofluorescent imaging was performed on 12 hepatocellular carcinoma (HCC) patient surgical resections reported by our group in (*36*). Representative 6-color image showing hepatic nerves (NCAM+) with markers for smooth muscle (αSMA+), endothelial cells (CD34+), bile ducts (panCK+, EpCAM+, NCAM+), and immune (CD45+) cells from LHCC35 biopsy. (**I**) Nearest neighbor profile across 12 HCC surgical samples in HCC tumor adjacent samples. Frequency of nearest neighbor: manual identification analyses conducted from identically-sized regions with nearest cellular neighbors determined at three locations along the nerve using an 18-color antibody panel (see Supplemental File 2). Upper rectangles reveal the relative average distance (μm) of nearest neighbor across cell types. Lower circles represent relative amount of nerves/biopsy region. Statistical significance determined by unpaired Student’s *t* test: \**P* < 0.05; \*\**P* < 0.01; ns = not significant. αSMA = α-smooth muscle actin, DC = dendritic cell, EpCAM = epithelial cellular adhesion molecule, LYVE1 = lymphatic vessel endothelial hyaluronan receptor 1, NCAM = neural cell adhesion molecule, panCK = pan-Cytokeratin, TREG = regulatory T cell, and UITC = unconventional and innate-like T cells.

While vagal activity has been implicated in systemic metabolic processes and angiogenesis (*10*, *30*), vagotomized mice showed no abnormalities in serum chemistry tests measuring transaminases and albumin liver function (fig. s1D). Hepatic vascular endothelial growth factor A (VEGFa) sera levels were also comparable in vagotomized and sham controls (fig. s1E).

HV mice exhibited significantly smaller tumors across multiple liver tumor models (Fig. 1B-D). Reduced tumor growth and burden were observed in female C57BL/6 HV mice following intrahepatic injection of syngeneic RIL175 (HCC) cells (Fig. 1C, D, and fig. s1F). As sex differences have been reported in liver cancer pathogenesis (*31*), we repeated and confirmed findings in male mice (fig. s1G). We also observed HV tumor reduction in established metastatic models (*32–34*). Compared to SV controls, HV livers developed smaller B16-F10 (melanoma) tumors within C57BL/6 mice (Fig. 1E and fig. s1H), while BALB/c animals exhibited fewer and smaller A20 (lymphoma) metastatic tumors following tail vein injection (Fig. 1F and fig. s1I). In addition, we performed subcutaneous tumor injection. Hepatic vagotomy failed to affect subcutaneous RIL175 tumor growth, highlighting liver-specific outcomes (Fig. 1G). In summary, hepatic vagotomy robustly controlled liver tumors independent of cell line, tumor model, and mouse sex/strain.

To explore a causal role for nerve activity in the tumor microenvironment within liver tumors, we analyzed differentially expressed genes (DEGs) from eight distinct human cancer tissues in publicly available datasets (TCGA [The Cancer Genome Atlas] and GTex [Genotype-Tissue Expression] (*35*). Our pathway enrichment analyses revealed significantly altered neurological pathways in seven of the eight cancer tissues, including liver (HCC), identifying increased neuroinflammation and neuronal projection within rim (tumor-adjacent) tissue (fig. s2A and Supplemental File 1). We also observed nerve infiltration within the HCC rim in a small clinical cohort (Fig. 1H and fig. s2B) recently characterized by our lab (*36*). A spatial immune atlas using CODEX (co-detection by indexing) revealed NCAM+ (neuronal cell adhesion molecule) nerves in 12 HCC surgical resections. We manually profiled nerve (> 50 µm branch, DAPI^-^ NCAM^+^) neighborhoods (Fig. 1I). Although precise neuronal identity cannot be determined by the 38-marker CODEX panel (see (*36*)), we identified specific immune, stromal, and biliary features (fig. s2C and Supplemental File 2). Adaptive immune cells (CD45+CD3+) comprised the largest percentage of nearest cellular neighbors at ∼42% averaged across HCC surgical specimens (Fig. 1I and fig. s2D). As nerves within the periphery regulate adaptive immune activity (*15*, *25*), we next assessed whether cholinergic neuroimmune interactions shaped HV outcomes.

### ACh activity regulates liver tumor burden via CD8+ T cells

Parasympathetic vagal nerves exert divergent outcomes in non-liver tumor models, either preventing or promoting tumor growth via ACh release (*19–23*). As expected, HV livers exhibited reduced ACh levels compared to sham controls (Fig. 2A). To specifically ascertain cholinergic effects, we treated mice with bethanechol (Fig. 2B), a broad, muscarinic AChR agonist that altered tumor cell proliferation in pancreatic cancer models (*22*).

**Figure 2.**
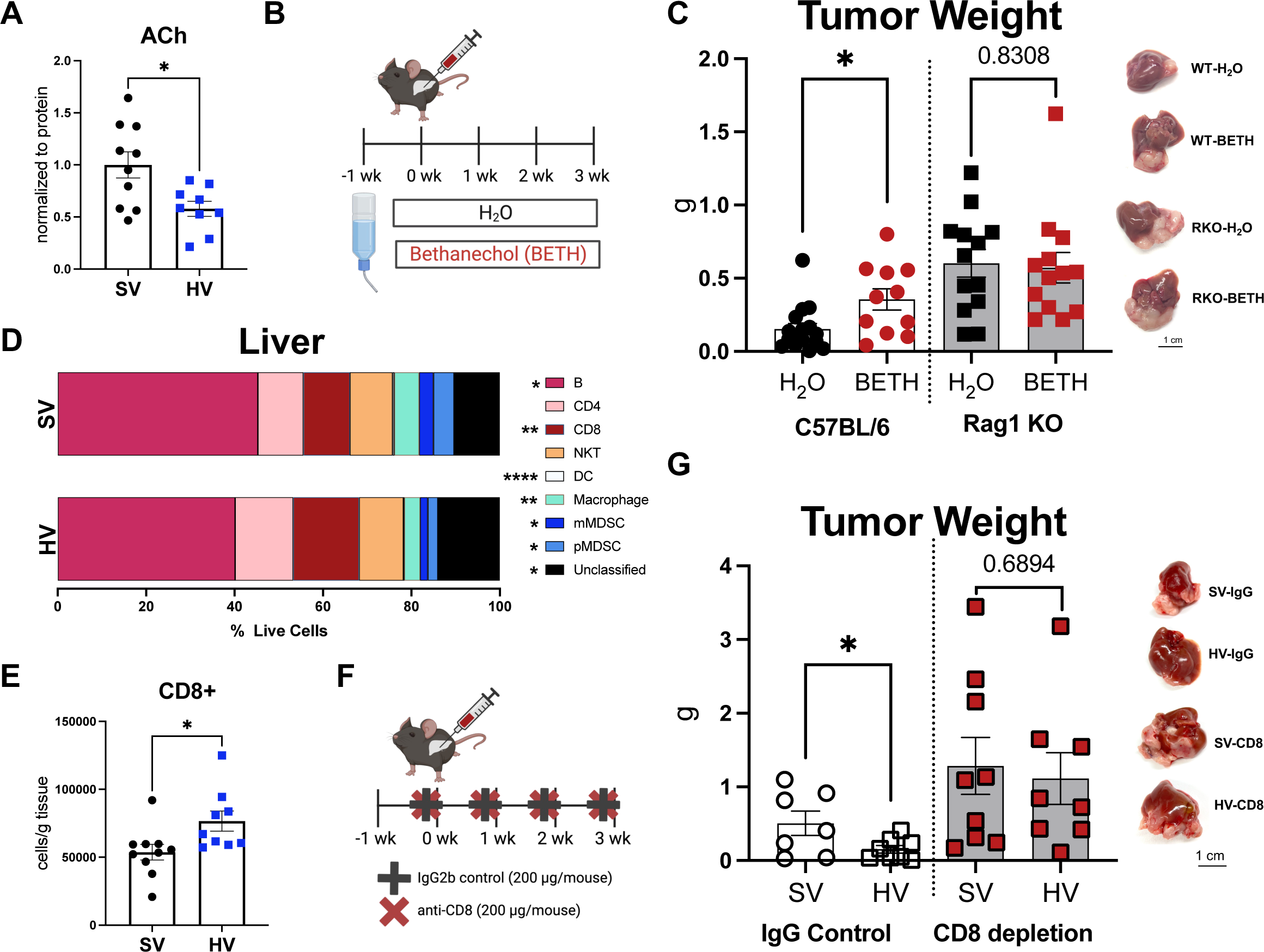
Hepatic vagotomy controls liver tumors via cholinergic signaling and CD8+ T cells. **(A)** Hepatic ACh levels measured via ELISA and normalized to tissue protein content. Data assessed 21 d following RIL175 tumor initiation. (**B**) Systemic bethanechol administration model: bethanechol (400 µg/ml drinking water) was provided 3-5 d prior to tumor initiation and tumor burden was assessed at 21 d following RIL175 intrahepatic injection. (**C**) Tumor weights and representative liver images from C57BL/6 (WT) and Rag1KO (RKO) bethanechol (BETH) models. Data pooled from two independent cohorts, *n* = 16 B6_H_2_O, 11 B6_BETH, 13 RKO_ H_2_O, 13 RKO_BETH. (**D**) Hepatic immune cell composition assessed via flow cytometry as average frequency (% live cells), *n* = 10 SV, 9 HV. (**E**) Cell counts of hepatic CD8+ T cells from **D** normalized to tissue weight. (**F**) CD8+ T cell antibody depletion tumor model: C57BL/7 mice were randomized and treated with IgG2b control or anti-CD8 (200 μg/mouse, 1x / wk starting 1 d before intrahepatic injection). (**G**) Tumor weights and representative liver images from RIL175 tumor weights at 21 d, *n* = 7 SV-IgG, 8 HV-IgG, 9 SV-aCD8, 8 HV-aCD8. Data in **A**, **C**, **G**, and **E** represent mean ± SEM. Statistical significance determined via unpaired Student’s *t* test: \**P* < 0.05; \*\**P* < 0.01; \*\*\**P* < 0.005; \*\*\**P* < 0.001; ns = not significant.

Systemic bethanechol provision (BETH: 400 μg/mL drinking water) increased HCC (RIL175) tumor burden (fig. s3A, B) without impacting water consumption (fig. s3C). Direct bethanechol exposure failed to promote RIL175 proliferation (fig. s3D), suggesting an indirect cholinergic mechanism. As ACh activity influences systemic inflammatory responses (*22*, *37*), we repeated HCC models in an independent C57BL/6 cohort alongside age-matched Rag1KO mice on a C57BL/6 background. While bethanechol promoted RIL175 tumor growth in C57BL/6 mice, bethanechol failed to alter tumor burden in Rag1KO mice lacking mature adaptive immunity (Fig. 2C). As Rag1KO-H_2_O and Rag1KO-BETH livers had comparable myeloid accumulation (fig. s3E), results indicated an ACh-adaptive immune mechanism.

We then profiled immune composition within vagotomized models to identify putative adaptive immune participants. Hepatic vagotomy significantly promoted anti-tumor immune profiles in liver and tumor specimens from HCC mouse models (Fig. 2D and fig. s4A, B). In contrast, splenic immune cell frequencies were largely comparable in HV and SV mice (fig. s4C), highlighting liver-specific cholinergic modulation.

Flow cytometry analyses revealed a decrease in B cells, dendritic cells (DCs), macrophages, and myeloid-derived suppressor cells (MDSCs), including monocytic (mMDSC) and granulocytic/polymorphonuclear (pMDSC) types within HV livers. In contrast, HV livers exhibited an expansion within CD3+ T cells (Fig. 2D), an HV feature maintained within metastatic B16-F10 and A20 livers (fig. s4D, E). While gut-liver interactions have been reported to control Treg (T regulatory cell) accumulation (*25*, *29*), brain-liver disruption did not alter hepatic Treg (CD4+Foxp3+) abundance by flow cytometry (fig. s4F). Only CD8+ T cells exhibited significant accumulation within HV livers, assessed as frequency and absolute count (Fig. 2D, E).

We next utilized CD8+ T cell antibody depletion to confirm CD8+ T cell involvement (fig. s4G, Fig. 2F). While vagotomy reduced HCC tumor burden within IgG isotype controls, CD8+ T cell depletion abrogated HV tumor control (Fig. 2G). Collectively, these results indicate that both muscarinic ACh signaling and CD8+ T cells are required for vagal-dependent tumor control in liver cancer. To establish whether HV tumor control influences specific CD8+ T cell subsets and identify precise cholinergic-CD8+ T cell interaction, we utilized transcriptional and *in vitro* studies.

### Cholinergic disruption promotes CD8+ T cell effector activity

We performed scRNA-Seq analyses on sorted (Live+CD45+) immune cells from three SV and three HV livers with matched (pooled) tumor samples derived from mice with orthotopic RIL175 HCC tumors (Supplemental File 3). Following stringent quality control, 32,722 cells were utilized for downstream analyses (5,980 median UMI averaged across samples). Thirteen subsets emerged during unsupervised clustering, including myeloid, B cell, NK, and ILC1 populations (Fig. 3A and fig. s5A-D) with scRNA-seq results largely reflecting earlier flow cytometry analyses (Fig. 2D). While HV mice exhibited a moderate expansion of hepatic NK and ILC1 populations (c12_NK and c13_ILC1, fig. s4C), characterized by distinct gene signatures reported in (*38*), the overall HCC immune transcriptome was largely characterized by distinct T cell (Live+CD45+CD3+) subclusters (Fig. 3A, fig. s5A-D, and Supplemental File 3).

**Figure 3.**
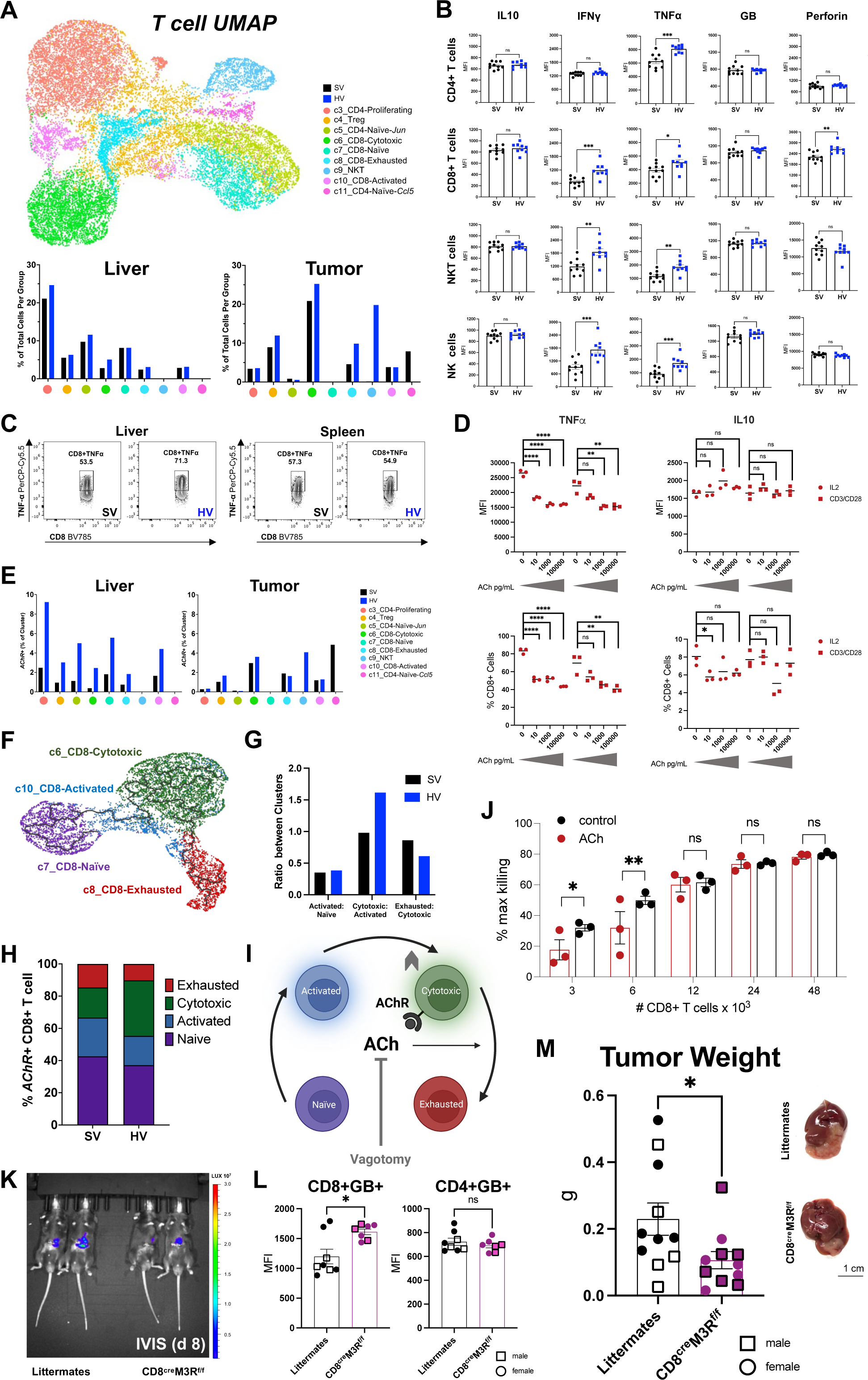
ACh signaling promotes CD8+ anti-tumor response. **(A)** (*Top*) T cell UMAP (Live+CD45+CD3+) from SV and HV RIL175 cohort for matching liver (*n* = 3 / group) and (pooled) tumor samples. (*Bottom*) frequency of total cells within scRNA-Seq unsupervised T cell clusters from liver (tumor adjacent) and tumor (tumor-infiltrating lymphocytes) samples. (**B**) *Ex vivo* intracellular cytokine and cytotoxic markers from hepatic lymphocytes as measured by flow cytometry median fluorescence intensity (MFI) provided. Liver samples collected 21 d following RIL175 tumor initiation and PMA/ionomycin stimulated for 4 h prior to staining. (**C**) Representative gating of CD8+TNFα+ population in liver (*left*) and spleen (*right*) from RIL175 tumor model. (**D**) Following expansion under IL2 (100 ng/mL) or CD3/CD28 Dynabeads (3:1 T cells: Dynabeads) stimulation for 48 h, splenic cells underwent ACh exposure (2 h) plus PMA/ionomycin activation + ACh (4 h). Flow cytometry of TNFα+ and IL10+ staining in CD8+ T cells (*n* = 3 / technical replicates), MFI reported. **(E)** The percent of positive *AChR* expression within scRNA-Seq lymphocyte clusters (Live+CD45+CD3+). (**F**) Monocle trajectory analyses of CD8+ T cell hepatic subclusters (*n* = 3 SV, 3 HV). (**G**) Ratio of *AChR*+ CD8+ T cell naïve and effector subclusters from SV and HV samples. (**H**) Frequency of CD8+ T cell subclusters in SV and HV samples. Data in **E**-**H** from biological samples reported in **A**. (**I**) Cholinergic anti-cytotoxicity CD8+ T cell model. ACh disruption (*e.g*., vagotomy) prevents AChR-dependent alteration of cytotoxic CD8+ T cell subsets (increased in HV samples), maintaining effector populations. (**J**) Cytotoxicity assay from splenic OT-1 CD8+ T cells after 12 h coculture with B16-F10-OVA tumor cells with increasing effector-to-tumor target ratios. Flow cytometry analyses of ACh (1 µg/ml) or media controls with maximum killing determined using heat-killed controls (*n* = 3 / technical replicates). (**K**) Representative IVIS® Spectrum in vivo imaging 8 d after RIL175 intrahepatic injection in CD8^cre^M3R^f/f^ mice and littermate controls. (**L**) CD8+GB+ in CD4+ and CD8+ T cells intracellular staining following 4 h PMA/ionomycin stimulation, MFI reported. (**M**) RIL175 tumor weights measured at 14 d, square = male mice, circle = female mice (*n* = 11 / group). Data pooled from two independent cohorts. Unless noted, statistical significance determined by unpaired Student’s *t* test or one-way ANOVA with *posthoc* Dunnett’s test (**C**) or two-way ANOVA (**J**): \**P* < 0.05; \*\**P* < 0.01; \*\*\**P* < 0.05; \*\*\*\**P* < 0.001; ns = not significant. Bar graphs and **D** display mean, reported whiskers represent ± SEM; GB = granzyme B.

After batch correction for liver and tumor samples together, UMAP (Uniform Manifold Approximation and Projection) revealed nine T cell subclusters with tissue- and vagal-specific profiles (Fig. 3A and fig. s5D-F). Cluster identity of liver- and tumor-infiltrating lymphocytes was defined by gene signatures and guided by prior scRNA-Seq landscape reports, fig. s5F and Supplemental File 3 (*36*, *39*). The largest CD4+ T cell subcluster, c3_CD4_Proliferating, was characterized by high expression of *Il2rb* and *Id2*, markers involved in T cell maintenance and proliferation (*40*, *41*). HV livers, but not HV tumors, displayed increased c3_CD4_Proliferation frequency. In contrast, the c4_Treg subsets (*Foxp3* high) were more prevalent within tumor samples. HV tumors displayed a higher frequency of the immunosuppressive c4_Treg subset, perhaps reflecting an overall increase of CD4+ T cells after vagotomy. These differences were less pronounced within liver samples, matching flow cytometry analyses (fig. s4F). Remaining CD4+ T cell subclusters, c5_CD4_Naïve-*Jun* and c11_CD4_Naïve-*Ccl5*, exhibit largely comparable gene signatures characterized by high *Ccr7*, *Lef1*, and *Sell* expression, matching previously reported cluster annotation (*39*). NKT cells (c9_NKT) were significantly increased within HV tumors. While hepatic vagotomy markedly altered the liver immune landscape, further transcriptional analyses focused on CD8+ T cell subsets.

CD8+ T cells with naïve transcriptional signatures (c7_CD8_ Naïve subcluster) were observed in HV and SV liver samples (Fig. 3A). The predominant CD8+ T cells subcluster, c6_CD8_Cytotoxic, exhibited high granzyme expression, specifically *Gzma* and *Gzmb,* gene signatures (fig. s5F and Supplemental File 3). Indeed, the c6_CD8_Cytotoxic cluster comprised the largest percentage (>25%) of the HV tumor CD45+ immune landscape. The c6_CD8_Exhausted subset, characterized by high expression of exhaustion marker *Tox* (thymocyte selection-associated high mobility group box), was also higher within HV mice. This subset exhibited moderate cytotoxic gene expression (*Gzma*, *Gzmb*). As both exhausted and late-differentiated effector memory CD8+ T cells express *Tox* (*42*), this subset likely contains a mix of effector and exhausted properties. The smallest CD8+ T cell cluster, c10_CD8_Activated was largely comparable across HV and SV samples and characterized by the highest expression of *Cd69,* a marker of early T cell activation (*36*). Collectively, hepatic vagotomy promoted anti-tumor adaptive features, characterized by elevated CD8+ T cell effector properties (Fig. 3A). While flow cytometry and transcriptional analyses provided functional profiling of the HV T cell compartment, subsequent studies validated functional activity.

To address effector function, liver-infiltrating lymphocytes from HCC-bearing mice underwent *ex vivo* stimulation (PMA/ionomycin) as reported (*36*). We observed increased frequency and mean fluorescence intensity (MFI) expression of proinflammatory cytokines IFNγ and TNFα across HV lymphocytes (Fig 3B, fig. s6A). In contrast, anti-inflammatory cytokine IL10 was unaltered by hepatic vagotomy. Reflecting transcriptional cytotoxic profiles, CD8+ T cells also showed increased perforin expression (Fig. 3B). *Ex vivo* analyses of matched splenic lymphocytes revealed comparable CD8+ T cell activation in SV and HV RIL175 models (Fig. 3C and fig. s6B), confirming organ-specific vagal modulation.

Metastatic liver models displayed comparable HV-induced inflammatory markers, (fig. s6C, D), highlighting a robust hepatic neuroimmune arc. While cholinergic disruption promoted anti-tumor immunity responses, we next sought to establish whether ACh activity directly influenced CD8+ T cells.

### ACh signaling affects CD8+ T cell anti-tumor activity

To assess direct ACh-CD8+ T cell interaction, we measured intracellular cytokine production following acute (4 h) ACh exposure as described (*27*, *36*). Splenic lymphocytes were expanded with IL2 or CD3/CD28 as reported (*43*). ACh exposure reduced CD8+ T cell production of TNFα, but not IL10, in a dose-dependent manner (Fig. 3D), reflecting prior reports of ACh dampening macrophage cytokine release (*27*, *28*). We observed similar cytokine modulation within ACh-treated CD4+ T cells, indicating broad lymphocyte control (fig. s7A).

We next examined whether muscarinic AChR engagement was directly involved in CD8+ T cell modulation. While ACh depletion via vagotomy promoted CD8+TNFα+ expression, bethanechol treatment reduced intracellular TNFα production in *ex vivo* splenic T cells (fig. s7B). In contrast, exposure to 4-DAMP (1,1-dimethyl-4-diphenylacetoxypiperidinium iodide), a selective AChR antagonist preferentially targeting CHRM3, promoted TNFα+ expression in a dose-dependent manner within stimulated CD8+ T cells (fig. s7C). While these findings do not preclude direct ACh-tumor activity, ACh exposure (48 h) failed to impact proliferation or viability of tumor cell lines in vitro (fig. s7D).

Hepatic vagotomy depletes liver ACh levels (Fig. 2A). Intriguingly, scRNA-seq analyses revealed increased *AChR+* subsets across most HV lymphocytes, suggesting a compensatory upregulation after vagotomy (Fig. 3E and fig. s7E). Subsequent pathway enrichment analyses identified subset-set specific cholinergic impact. To focus on ACh-immune activity, we developed a “Neuroimmune Panel” (Supplemental File 4) including gene ontology (GO) pathways reporting cholinergic activity and pro-inflammatory activation expressed in our hepatic tissue. As anticipated, HV liver and tumors (Live+CD45+) cells expressed enriched proinflammatory responses, including Lymphocyte Activation Involved in Immune Response (GO:0002285) and Neuroinflammatory Response (GO:0150076), fig. s8A, B and Supplemental File 4.

Although all HV CD8+ T cell subsets exhibited enriched neuroinflammatory expression (fig. s8B), ACh response (Cellular Response to Acetylcholine [GO:1905145] varied, fig. s8C. In contrast with remaining CD8+ T cell subsets, only the c10_CD8_Activated subset exhibited enriched ACh cellular responses in vagotomized mice. In contrast, more abundant c6_CD8_Cytotoxic and c8_CD8_Exhausted subsets exhibited elevated ACh cellular responses within SV samples, suggesting that ACh exhibits subset-specific interactions.

This proposal is supported by time-series analyses of liver and tumor CD8+ T cells. Pseudotime and trajectory analyses using Monocle3 (*44*) revealed dynamic CD8+ T cell branching across trajectory pathways (Fig. 3F). Only the c6_CD8_Cytotoxic subset contained trajectory branching into the exhausted cluster. Indeed, ratios of *AChR+* T cell clusters in liver and tumor samples revealed comparable activated: naïve subset ratios in SV and HV mice, while HV samples had elevated cyotoxic:activated subsets and reduced exhausted: cytotoxic subsets (Fig. 3G), frequency patterns reflected in *AChR+* CD8+ T cells (Fig. 3H). Collectively, AChR agonist models (Fig. 2B), in vitro studies (Fig. 3D), and transcriptomic analyses (Fig. 3E-H and fig. s8) suggest that disruption of ACh (*e.g.*, via vagotomy) blocks AChR signaling, facilitating maintenance and expansion of CD8+ T cell cytotoxic features via AChR signaling (Fig. 3I).

To validate ACh modulation of cytotoxicity within a tumor setting, we isolated CD8+ T cells from transgenic OT-1 mice that express CD8+ T cells specific for an MHC-I class-restricted, ovalbumin-derived peptide; and cocultured OT-1 CD8+ T cells with B16-F10-OVA tumor cells. ACh administration impaired CD8+ T cells lysis of B16-F10-OVA cells (Fig. 3J).

ACh robustly suppresses CD8+ T cell effector function via muscarinic AChR-dependent signaling. Antagonist studies (fig. s7C) and transcriptional analyses (fig. s9A) suggest that the muscarinic CHRM3 receptor, a regulator of peripheral tumor cell proliferation (*21*, *22*, *45*), contributes to the modulation of CD8+ T cell cytotoxic function. Consequently, we utilized Cre/*lox* manipulation to generate mice that lacked *Chrm3* expression on CD8+ T cells (CD8^cre^M3R^f/f^). Compared to littermate controls, CD8^cre^M3R^f/f^ mice exhibited a modest reduction in HCC tumor growth (Fig. 3K and fig. s9B) in non-vagotomized mice. Notably, CD8^cre^M3R^f/f^ hepatic CD8+ T cells, but not CD4+ T cells, exhibited increased granzyme B expression (Fig. 3L), a marker of cytotoxic activity. While CD8^cre^M3R^f/f^ mice displayed significantly smaller tumor burden compared to littermates (Fig. 3M), genetic modulation did not impair body weight, non-tumor liver weight, or alter total lymphocyte accumulation (fig. s9C, D), further supporting targeted modulation of CD8+ T cell function.

In summary, this work reveals a dynamic vagal ➔ liver axis. Hepatic ACh activity elicits diverse and robust modulation of liver CD8+ T cell subsets, specifically affecting cytotoxic responses via CHRM3 signaling (Fig. 3). Beyond cholinergic alteration, liver cancer disrupts neurological-associated pathways in mice (fig. s8) and men (fig. s2A). Interorgan crosstalk is inherently bidirectional (*15*, *16*) and vagal ➔ brain arcs likely shape neurological side effects linked to HCC. We next explored whether hepatic vagotomy altered behaviors associated with liver cancer (*e.g*., aberrant fatigue) (*46–49*).

### Vagotomy benefits neurological function in HCC tumor-bearing mice

Following facility acclimatization, vagotomized HCC mice underwent a battery of established behavioral tests, including phenotyping, open field test (OFT), and Y-maze (Fig. 4A) (*48*, *50*, *51*). To address neurological behaviors elicited by vagotomy versus tumor, a “Baseline” series of tests were performed prior to RIL175 tumor implantation. The latter timepoints assessed behaviors during periods when SV and HV mice exhibited smaller and comparable tumor burden (Early), as well as larger tumors and significant tumor variation (Late), based on prior *in vivo* imaging (Fig. 1C). While exercise decreased liver tumor burden via a previously reported sympathetic-innate immune mechanism (*52*), modest locomotive tests did not impact HV tumor control (fig. s10A).

**Figure 4.**
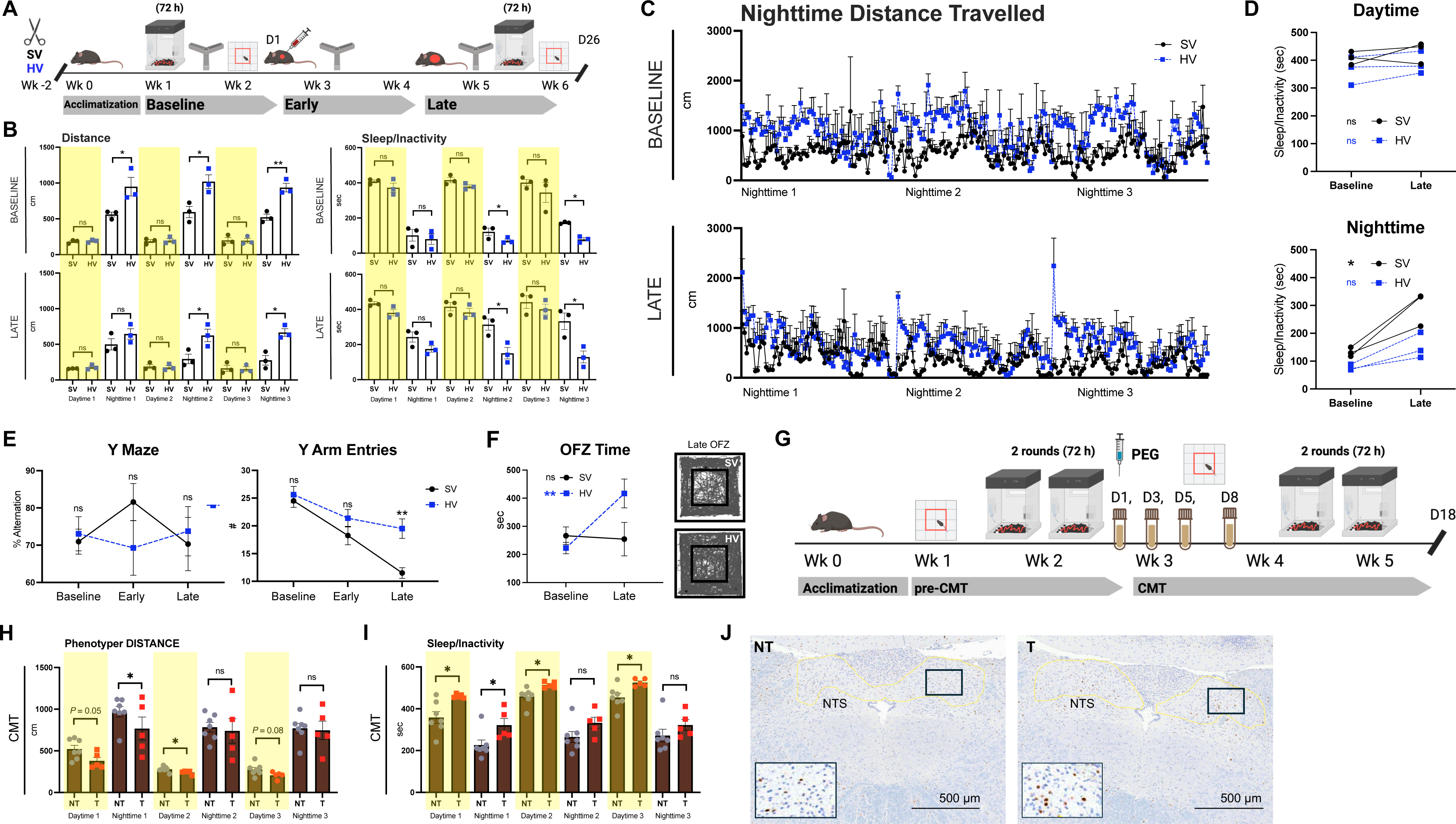
Impaired vagal signaling shapes neurocognitive response. (**A**) Schematic of behavioral analyses including facility acclimatization, Baseline (post-vagotomy, tumor free), Early (post-tumor), and Late (post-tumor) timepoints with 72-h phenotyper tracking, Y-maze, and open field test (OFT) testing; *n* = 8 SV, 8 HV. (**B**) (*Left*) phenotyper distance (total distance moved from center-point) averaged across 12 h at Baseline (*top*) and Late (*bottom*) timepoints. (*Right*) immobility (sleep/inactivity) tracking at Baseline (*top*) and Late (*bottom*), 12-h timepoints. (**C**) Phenotyper nighttime distance, points indicate 10-min intervals, at Baseline (*top*) and Late (*bottom*) timepoints; *n* = 3 SV, 3 HV in phenotyping experiments. Availability of PhenoTyper 3000 cages limited phenotyping, remaining mice were placed in single housing throughout phenotyping experiments. (**D**) Immobility in SV and HV mice averaged across 72-h tracking, paired baseline and late timepoints. (**E**) Y maze percent alternation (*left*) and total arm entries (*right*) at Baseline, Early, and Late timepoints (5 min free exploration). (**F**) (*Left*) total time spent in open field zone (OFZ) in paired Baseline and Late timepoints (30 min free exploration). (*Right*) representative OFT tracking plots for SV and HV late timepoint, *n* = 8 SV, 8 HV. Mice in **B**-**F** from the same cohort. (**G**) Non-tumor (NT) and tumor (T) cecal microbiota transplant (CMT) recipients and behavioral analyses timeline: T donors received intrahepatic injection of RIL175 cells in PBS/Matrigel matrix, while NT donors underwent intrahepatic injection of PBS/Matrigel matrix vehicle. Cecal content collected at 21 d and pooled, filtered content was transplanted into NT or T recipients via four rounds of oral gavage (100 μL) after polyethylene glycol (PEG) laxative treatment. Phenotyper and OFT testing performed at Baseline (pre-cecal microbiota transplant = pre-CMT) and CMT timepoints. (**H**) Phenotyper mobility (total distance moved from center-point) during CMT testing and averaged across 12 h (*n* = 7 NT, 5 T). (**I**) Daytime immobility measured from mice averaged across 12 h. Highlighted overlay for all phenotyper data indicative of daytime hours. (**J**) Representative images of the brain stem from RIL175 tumor-bearing mice (RIL175) or non-tumor controls (CON) that underwent intrahepatic injection, NTS highlighted. Statistical significance determined by unpaired or paired Student’s *t* test: (reported value) 0.05<*P*<0.1, \**P* < 0.05; \*\**P* < 0.01; ns = not significant or reported value provided. Whiskers reflect ±SEM and bar graphs indicate mean; NTS = nucleus tractus solitarius.

We first assessed fatigue in mice housed in PhenoTyper 3000 cages, capturing daytime (rest) and nighttime (mobile) activity and immobility (*50*). Vagotomized mice exhibited decreased fatigue, as measured by mobility (distance traveled) and immobility (sleep/inactivity), specifically during nighttime cycles (Fig. 4B-D and fig. s10B). While liver cancer exacerbated exhaustion in SV and HV mice, HV mice maintained shorter sleep/inactivity periods (Fig. 4B-D).

Beyond locomotion, we assessed cognitive function and anxiety-like behaviors, via the Y maze and OFT (*48*, *51*), respectively. In each Y maze assessment, SV and HV mice displayed normal spontaneous alternation (Fig. 4E), suggesting comparable cognitive ability regarding exploration and acute spatial memory (*48*). At the Late timepoint, HV animals exhibited increased mobility assessed by total Y maze arm entries, as well as total distance travelled, average speed, and immobility (Fig. 4E and fig. s10C). Increased mobility was also observed in Late, but not Baseline, OFT trials (fig. s10D). Upon liver tumor development, HV mice showed reduced anxiety-like behavior, demonstrated by increased open field zone (OFZ) exploration (Fig. 4F and fig. s10D). In summary, tumor-bearing HV mice exhibit improved neurological outcomes, notably reduced fatigue, increased mobility, and better anxiety-associated behavior.

While behavior potentially reflects different SV and HV tumor burden, final tumor weight did not correlate with either Y maze or OFT activity (fig. s10E). While hepatic vagal integrity shaped liver tumor burden (Fig. 1D-F), SV tumor size was not a significant driver of behavioral impairment (fig. s10E). Consequently, we hypothesized that systemic HCC dysbiosis shapes neurological outcomes. Intriguingly, non-hepatic vagal modulation of behavior has been reported in the context of gut-vagal-brain interaction (*17*). Largely comprised of afferent fibers, vagal nerves encompass the gastrointestinal (GI) tract and respond to microbial cues (*17*, *53*). Consequently, we explored whether the gut microbiota from HCC mice was sufficient to shape cancer-associated behavior.

We performed cecal microbiota transplant (CMT) via oral gavage from pooled RIL175 tumor-bearing donors (T) or intrahepatic surgical controls without tumors (NT), see Fig. 4G. Phenotyping and OFT trials were performed prior and following CMT (Fig. 4H). While T and NT recipients exhibited comparable OFT exploration and activity (fig. s11A, B), phenotyping revealed reduced mobility (distance traveled) and increased immobility (sleep/inactivity) within T recipients post CMT (Fig. 4H, I and fig. s11C-E), suggesting that gut microbiota dysbiosis from HCC livers is sufficient to promote cancer-associated fatigue. In contrast to earlier behavior analyses in vagotomized mice (Fig. 4B-D), the CMT study reported altered exhaustion behaviors primarily during the daytime (resting period: Fig. 4I), likely linked to circadian disruption from repetitive CMT.

These results indicated that gut microbes are sufficient to alter cancer-associated behavior. As noted, the GI tract is heavily innervated by vagal afferents, sensory fibers that relay information into the nucleus tractus solitarius (NTS) within the brainstem (*29*). Compared to a non-tumor bearing controls, HCC mice exhibit modest NTS activation as defined by c-Fos expression (Fig. 4J and fig. s11F). These findings indicate that cancer-associated gut dysbiosis influences brain features. Both vagotomized mice (Baseline) and CMT recipients, however, exhibit altered fatigue patterns (Fig. 4B-D). Consequently, we investigated whether pertinent gut microbiota dysbiosis primarily resulted from vagotomy and/or liver cancer.

### Tumor, not vagotomy, robustly shapes fecal microbiota composition

Vagal activity influences GI transit time, which itself is a major driver of gut microbiota composition (*54*). Truncal (subdiaphragmatic) vagotomy impairs gastric emptying and GI transit, requiring an accompanying pyloroplasty (*24*, *45*, *55*). Precise hepatic branch vagotomy, however, failed to alter GI transit time (fig. s12A). Indeed, cohoused non-tumor bearing vagotomized (HVx) and sham controls (SVx) exhibit comparable fecal gut microbiota composition and alpha diversity assessed by 16S rRNA-Seq (fig. s12B, C). Indeed, cage effect, rather than vagotomy status, had greater influence on gut microbiome profiles (fig. s12B and Supplemental File 5).

In contrast to immune assessments in tumor-bearing mice (Fig. 2D and Fig. 3A), SVx and HVx livers exhibited similar immune cell compositions (fig. s12D, E) and comparable pro-inflammatory cytokine expression in CD4+ and CD8+ T cells (fig. s12F, G). These findings indicate that the hepatic tumor is required for vagotomy-dependent immune alteration. We next utilized cohousing experiments to further establish the role of hepatic vagotomy-induced microbiota dysbiosis and liver tumor burden.

Coprophagic activity and reciprocal grooming in mice normalizes the fecal microbiota composition during prolonged cohousing (*56*). Following vagotomy procedure, a subset of randomized SV and HV mice were cohoused (cSV and cHV, respectively) prior to intrahepatic injection of RIL175 cells (fig. s12H). Cohousing failed to alter tumor-bearing GI transit time (fig. s12I), with transit times comparable to non-tumor bearing results (fig. s12A). All groups exhibited similar fecal microbiota alpha diversity with the co-housed beta diversity (Jaccard) and composition (Order) displaying an intermediate phenotype between the SV and HV fecal microbiome (fig. s12J-L and Supplemental File 5). While cohousing normalized gut microbiota profiles, cohoused mice had comparable tumor burden, liver weights, and CD8+TNFα+ expression as non-cohoused counterparts (fig. s12M-O). While non-cohoused SV and HV mice exhibited distinct fecal microbiota composition indicating vagal-dependent dysbiosis (fig. s12J-L), liver cancer markedly altered gut microbiota composition (fig. s12P). Post-tumor SV and HV gut microbiota features were more similar than fecal microbiota samples collected pre-tumor initiation (fig. s12P-R and Supplemental File 5), indicating that HCC, rather than vagotomy, has a stronger impact on the fecal gut microbiota composition. Collectively, cohousing studies indicate that vagal-induced dysbiosis of the gut microbiota was not sufficient to regulate hepatic CD8+ T cell anti-tumor immunity and subsequent liver tumor burden.

### Gut microbes modulate liver immunity via the vagus nerve

Our lab previously reported that gut bacteria modulate liver tumor burden via a gut-liver arc influencing hepatic NKT accumulation (*33*, *57*). Cohousing experiments (fig. s12) suggested that “direct” vagotomized liver-gut interactions do not impact CD8+ T cell accumulation and anti-tumor function. Behavioral CMT studies (Fig. 4H, I), however, suggested that liver cancer effects can be mediated via “indirect” gut microbiota-brain signaling. Thus, we hypothesized that the gut microbiota helps maintain hepatic immune tolerance via the vagus nerve, vagal disruption (e.g., vagotomy) facilitates expansion of anti-tumor immunity.

To specifically assess HCC-dependent microbiota dysbiosis, we performed CMT from eight RIL-175 tumor-bearing donors (T) and seven non-tumor bearing controls (NT), Fig. 5A. Recipients underwent a sham or hepatic vagotomy prior to repeated oral gavage containing donor-specific cecal content. Both hepatic immune and gut microbiota profiling were conducted, with fecal samples collected prior to CMT and at experimental endpoint. In addition, small intestine (jejunal, ileal) and cecal content was collected to examine gut microbiota composition across the GI tract (Fig. 5B).

**Figure 5.**
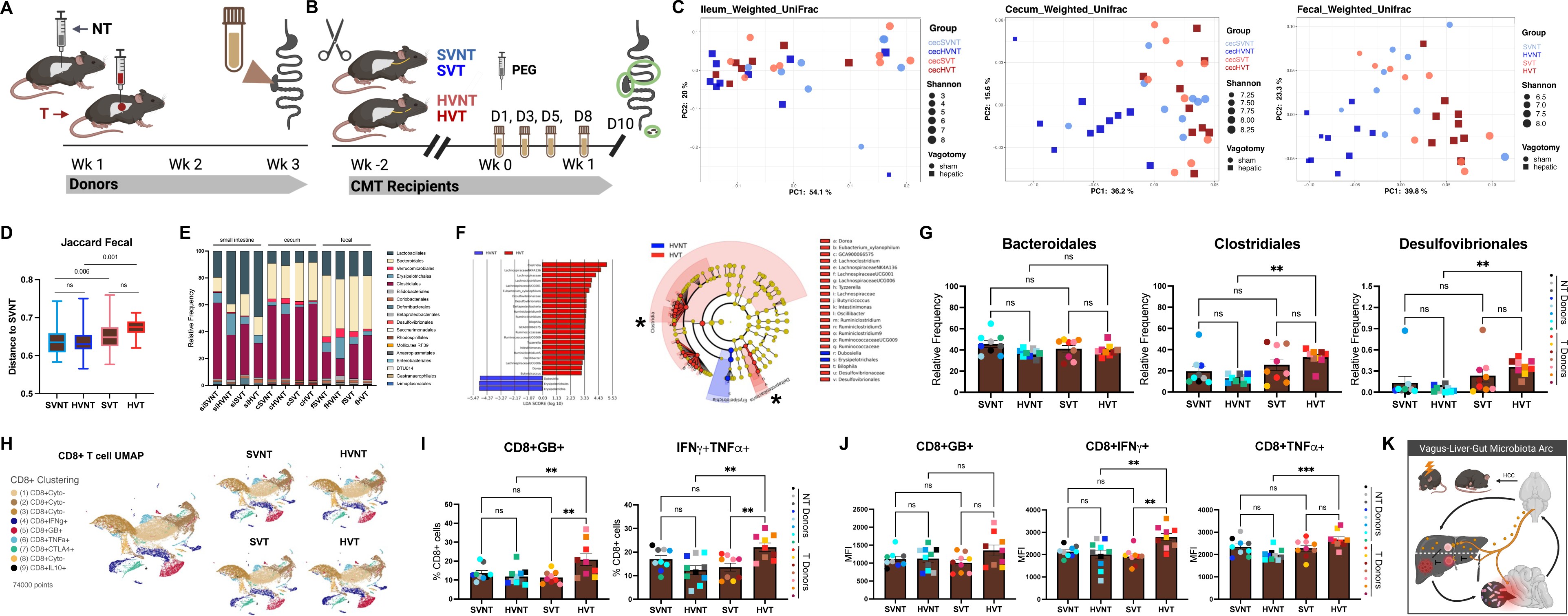
Gut microbes regulate liver immunity via vagus-liver arc. (**A**) Model of cecal microbiota transplant (CMT) recipient timeline: indicating vagotomy status (sham vagotomy = SV or hepatic vagotomy = HV) and unique donor (non-tumor = NT or RIL175 tumor = T). (**B**) Recipients received unique donor microbiota content. CMT procedure detailed in Expanded Methodology. Donors: *n* = 8 NT, 7 T; Recipients: *n* = 9 SVNT, 10 HVNT, 9 SVT, and 9 HVT. Green circles indicate collection region for recipient microbiota analyses via 16S rRNA-Seq 10 d following initial microbiota gavage. (**C**) Weighted UniFrac (beta diversity) analyses from CMT recipients assessing small intestinal, cecal, and fecal samples. Color = donor group, shape = vagotomy status, size = Shannon entropy (alpha diversity). (**D**) Jaccard distances (beta diversity of group) from fecal samples analyzed in **C**, pairwise PERMANOVA with q-value reported. (**E**) Fecal microbiota composition (Order) determined by 16S rRNA-Seq. (**F**) LefSE analyses (*84*) of HVNT and HVT fecal microbiota samples (Kruskal-Wallis α = 0.001, Wilcoxon test α = 0.001) with differentially expressed taxa determined via linear discriminant analyses (LDA) score >2 (*left*) and visualized via cladogram (*right*), asterisks identify two of the bacterial frequencies reported in **G**. (**G**) Relative abundance of fecal bacterial communities from CMT recipients, taxa represents Order. (**H**) UMAPs from Live+CD8+ hepatic cells analyzed by flow cytometry with fluorescence-based clustering identified via Partek Flow, (*left*) UMAP of all CD8+ T cells and (*right*) UMAPs separated into recipient group. (**I**) *Ex vivo* intracellular cytokine staining measured by manual gating of flow cytometry. Leukocyte stimulation (PMA/ionomycin) and staining as reported in Fig. 3B and Expanded Methodology. Percent of CD8+ T cells for granzyme B+ (GB+), IFNγ+TNFα+ subsets. (**J**) (*Left*-*right*) MFI of GB+, IFNγ+, and TNFα+ CD8+ T cells. Colors in **I** and **J** indicative of unique donor, while shape denotes vagotomy status. (**K**) Graphical representation of vagal-liver-gut arc regulating liver immunity. Liver cancer alters the gut microbiota and subsequent dysbiosis promotes (1) cancer-associated fatigue and (2) anti-tumor immune activity only within hepatic vagotomized mice. Vagal cholinergic signaling modulates the liver immune cell composition. Disruption of vagal activation facilitates anti-tumor immunity, notably via muscarinic signaling in CD8+ T cells. Expanded CD8+ T cell effector responses, in turn, controls liver tumor burden. Bar graphs indicate Tukey whisker plots (**D**) or mean ±SEM (**G**, **I**, and **J**). Unless noted, statistical significance determined by pairwise PERMANOVA (**D**) or via one-way ANOVA with *posthoc* Tukey’s multiple comparisons (**G**, **I**, and **J**): \*\**P* < 0.01; \*\*\**P* < 0.005; reported values or ns = not significant.

CMT from NT and T bacterial cocktails (1:1 cecal content: 50% glycerol) significantly altered the fecal microbiota composition of recipients (fig. s13A-C and Supplemental 5). The small intestine, cecum, and fecal samples displayed distinct, site-specific gut microbiota composition and diversity. The small intestine exhibited the lowest alpha diversity (Shannon entropy) with vagotomy status, not tumor, exerting a greater influence on bacterial composition (Fig. 5C, E, fig. s13D, and Supplemental File 5). In contrast, cecal and fecal bacterial composition reflected tumor (donor) status (Fig. 5C-E, fig. s13D, and Supplemental File 5), with fecal Jaccard distances (beta diversity) highlighting tumor-dependent alteration (Fig. 5D). Consequently, subsequent analyses focused on characterization of HCC tumor-dependent alterations within vagotomized mice (*i.e.,* HVNT vs HVT). Stringent LefSe (linear discriminant analyses [LDA] effect size: Kruskal-Wallis α = 0.001, Wilcoxon α = 0.001, LDA > 2) revealed numerous bacterial changes between the fecal HVNT and HVT microbiota (Fig. 5F), but no significant taxa enrichment in HVT versus SVT analyses, supporting tumor-dependent modulation. While LefSe did not identify any enriched taxa within the small intestine HVT samples, both cecal and fecal HVT communities exhibited increased Clostridial species (Fig. 5F, G and fig. s13E). Both HVT and SVT recipients exhibited increased expansion of Clostridiales, but not Bacteroidales, within the fecal microbiota, bacterial Order reported (Fig. 5G). The HVT fecal microbiota was also enriched in Desulfovibrionales (Fig. 5F, G), a sulfate-reducing taxon associated with liver malignancies and intestinal vagal stimulation (*58*, *59*). Bacterial commensals within Desulfovibrionales included *Bilophila* and *Desulfovibrio* (fig. s13F).

Microbiota transplantation has been linked to systemic immune modulation and altered CD8+ T cell activity (*60*). We subsequently profiled hepatic immune features to assess whether, and to what extent, the HCC (T) microbiota shapes CD8+ T cell accumulation and/or function. Comparable hepatic activation in SVT and HVT livers would suggest direct gut-liver interaction driven by HCC gut microbiota dysbiosis. In contrast, a vagus-liver model would exhibit distinct SVT and HVT responses, as SVT mice (intact vagus) would maintain cholinergic immune suppression, while HVT mice (vagal disruption) would exhibit elevated CD8+ T cell cytokine and cytotoxic function as observed in HV HCC models (fig. s14A and Fig. 3).

CMT recipients exhibited comparable body and liver weights (fig. s14B). Immune profiling via flow cytometry identified alteration in hepatic immune cell composition, notably a reduction in hepatic mMDSC cells in T recipients (SVT, HVT) compared to SVNT and HVNT counterparts. In contrast, CD8+ T cell accumulation was largely comparable across CMT groups (fig. s14C, D). We next assessed intracellular cytokine expression in PMA/ionomycin-stimulated hepatic lymphocytes. Partially supervised clustering of Live+CD8+ T cells revealed distinct cytokine and cytokine negative (Cyto-) subsets based on MFI expression (Fig. 5H). After parsing the CD8+ T cell UMAP by group, we observed comparable landscapes across SVNT, HVNT, and SVT profiles. The HVT UMAP was characterized by a relative decrease in Cyto-subsets and expansion c4_CD8_IFNγ+ and c5_CD8_GB+ clusters (Fig. 5H). Manual gating of flow cytometry datasets revealed elevated cytotoxic (granzyme B+) and inflammatory expression (IFNγ+TNFα+) across hepatic lymphocytes, particularly within CD8+ T cells (Fig. 5I, J and fig. s14E-G).

To summarize, microbiota transplantation from HCC donors (tumor-bearing mice) alters the fecal microbiota and, within HVT recipients, elicits hepatic anti-tumor activity. While both SVT and HVT mice received cecal transplantation from identical donors, HVT livers exhibited increased CD8+ T cell effector function, comparable with anti-tumor responses within HV liver cancer models (Fig. 3). While these findings do not preclude direct gut-liver activation, RT-qPCR of epithelial tissue revealed comparable inflammatory and barrier markers, including various claudin and tight junction genes (*Cldn2*, *Cldn4*, *Tjp1*), suggesting that HVT responses are not driven by leaky gut and/or intestinal inflammation from CMT (fig. s14H).

We report that cholinergic vagal disruption facilitates anti-tumor immunity, promoting CD8+ T cell cytotoxicity in liver cancer via a vagal-liver arc shaped by the gut microbiota. While the gut microbiome influences hepatic immune composition, function, and immunosuppression (*33*, *60*–*63*), ongoing research has largely studied direct gut microbiota-liver interactions. Our collective findings suggest that specific bacteria maintain hepatic tolerance via vagal activity.

Liver cancer promotes extensive intestinal dysbiosis (*64*). Intriguingly, T recipients exhibited increased relative abundance of sulfate-reducing bacteria previously reported to promote GI vagal activation (Fig. 5G and fig. s13F), (*17*, *51*). Cholinergic vagal activity maintains liver immune tolerance dampening pro-inflammatory and cytotoxic T cell responses. Downstream disruption of vagal activity (*e.g.*, via hepatic vagotomy), facilitates anti-tumor immunity via muscarinic AChR signaling, notably promoting CD8+ T cell effector responses. This work highlights an understudied cholinergic arc suppressing hepatic anti-tumor (Fig. 5K). We anticipate that further research may reveal microbial and neuroimmune therapeutic targets supporting liver cancer treatment.

## Discussion

Liver cancer is one of the deadliest cancers (*1*, *2*). Despite clinical advances, established immunotherapies largely fail patients due to poor immune responses (*9*, *65*). Surgical resection or ablative procedures remain the only curative treatments for liver cancers (*65*). Novel approaches, including cancer neuroscience efforts, are needed to promote hepatic anti-tumor immunity.

Interorgan crosstalk regulates immune responses (*15*). Within hepatic malignancies, interorgan discoveries have primarily focused on the gut-liver axis (*9*, *66*). Indeed, modulation of the gut microbiota revealed promising therapeutic modality in metabolic liver disease (*66–69*).

In cancer, gut microbes and microbial metabolites were first assessed as diagnostic biomarkers (*62*, *63*, *70*). Gut microbiota dysbiosis has been widely reported across liver cancers with specific taxa correlated to progression stages (*71*, *72*). Landmark studies revealed that the gut microbiome altered melanoma cancer outcomes and immunotherapy efficacy via inflammatory modulation (*73*, *74*, *61*). In HCC models, antibiotic exposure increased NKT cell accumulation, suppressing hepatic tumors in mice (*33*, *57*). These findings reveal dynamic interorgan modulation of liver anti-tumor immune responses.

We propose that gut-brain signaling contributes to hepatic immune tolerance via ACh activity. Precise hepatic vagotomy disrupts cholinergic-immune interactions, suppressing tumor growth. While HV mice exhibited broad immune alteration, including NKT expansion and MDSC control (Fig. 3A), Rag1KO studies and CD8+ T cell depletion studies (Fig. 2) highlighted a critical role for CD8+ T cell response. Although the extent of vagal innervation within healthy livers remains uncertain (*75*), cholinergic nerve fibers have been reported in liver malignancies and clinical liver cancer (*26*, *76*, *77*). Cholinergic signaling, notably via CHRM3, has been previously linked to gastric cancer proliferation via modulation of transcriptional pathways (*21*, *45*). Collectively, our work reveals that ACh inhibits CD8+ T cell effector function and identifies a CHRM3-dependent ACh-CD8+ T cell axis.

Recently, Zheng et al., 2023 (*25*) reported that intrinsic loss of cholinergic signaling in CD4+ T cells (*Chat* knockout) exacerbates liver cancer and promotes Treg expansion. In contrast, vagotomy failed to influence Treg accumulation (fig. s3B) and controlled hepatic tumor burden (Fig. 1D). We show that ACh activity exerts subset-specific responses in CD8+ T cells, which likely explains disparate cancer outcomes. While vagal disruption of ACh activity maintained effector CD8+ responses (Fig. 2E), genetic *Chat* ablation within CD4+ T cells inhibited auto- and paracrine ACh-signaling, triggering T cell dysfunction and subsequently promoting hepatic tumor progression (*25*).

Beyond tumor trajectories, vagal disruption improved fatigue, a neurological symptom linked to tumor-dependent microbiota dysbiosis. Indeed, the microbiota from HCC-bearing mice was sufficient to provoke anti-tumor immunity features only within vagotomized mice (Fig. 5K). Microbiota transplantation from HCC donors altered gut microbiota profiles, notably hydrogen sulfide-producing taxa (*i.e.*, *Desulfovibrio*, *Bilophila*) linked to colorectal and liver cancer (*78*, *79*). Recently, microbial-derived hydrogen sulfide was reported to activate vagal afferents via modulation of calcium-selective vagal receptors (*58*), highlighting a possible mechanism for HCC-induced vagal-brain signaling. While alteration of the gut microbiome influences vagal activity (*17*), the precise pathways by which the intestinal microbiome stimulates vagal afferents remains poorly characterized and warrants future study. As vagal afferents do not penetrate the intestinal epithelium, microbial modulation likely occurs via complex signaling mechanisms, not linked to specific bacterial members. Potential mechanisms include vagal alteration of small intestinal microbiota communities, intestinal bacterial metabolites, bacterially synthesized neurotransmitters, microbial-induced inflammation, and/or bacterial modulation of vagal-enteroendocrine cell interaction (*53*, *80*). Hepatic branch vagotomy partially reduces innervation of the upper (duodenal) intestinal tract due to loss of gastroduodenal fibers that ramify from the common hepatic branch. Indeed, hepatic vagal anatomy likely explains why vagotomy exerts a stronger impact on the small intestinal microbiota compared to cecal or fecal communities.

Finally, we show that gut microbiota-brain signaling maintains liver immune tolerance via vagal activity. Collectively, these findings highlight a targetable ACh-CD8+ T cell axis and identify dynamic vagal-liver signaling. Harnessing microbiota-neuroimmune crosstalk provides a therapeutic strategy to improve immunotherapy efficacy with implications for personalized medicine and liver cancer outcomes.

## Methods Summary

Detailed methodology and research material are provided within Supplementary text (Expanded Methodology). Briefly, we performed surgical vagotomy, shearing the hepatic vagal branch where the nerve diverges from the left subdiaphragmatic vagal trunk, or a sham procedure to maintain vagal integrity, as reported in (*29*). Following recovery, a subset of mice underwent intrahepatic injection or tail vein injection of RIL175, B16-F10, or A20 cancer cell lines, as previously described (*32–34*, *81*). Our lab performed high throughput flow cytometry or scRNA-seq from liver, tumor, and spleen immune samples utilizing immune panels developed within the Greten Lab and in collaboration with the NCI Flow Cytometry Core and Single-Cell Analysis Facility. Downstream analyses and visualizations were completed with FlowJo (vs 10+), Partek Flow, RStudio (R vs 4.2.1+), and Prism (vs 9+). Experimental designs, bioinformatic analyses, and data availability further detailed within Expanded Methodology.

Experimental assessment of lymphocyte cholinergic activity was performed via flow cytometry, ProcartaPlex immunoassays (Luminex®), ELISA, RT-qPCR, MTT Assay, and/or cytotoxicity/intracellular cytokine assays utilizing ACh, bethanechol, and 4-DAMP. *In vitro* findings were supported by *in vivo* studies in bethanechol-tumor models based on reported methodology (*22*), CD8+ T cell depletion studies guided by (*34*), and *Cre*/lox breeding strategies with Chrm3^f/f^ mouse mating pairs kindly provided by Jürgen Wess (NIH-National Institute of Diabetes and Digestive and Kidney Diseases [NIDDK]). Thorough experimental methodology for in vitro studies and animal tumor models listed within Expanded Methodology.

For systems biology studies, we conducted neurological and microbiome research. Behavioral tests included 72-h phenotyping (PhenoTyper, Noldus), Y-maze, and open field testing and were conducted by the NIH-NHLBI Murine Phenotyping Core. Histopathology was performed with the Molecular Histopathology Laboratory at Frederick National Laboratory and included standard H&Estaining (liver) and cFOS neuronal staining of the brainstem, guided by methodology reported in (*29*). Microbiota donors were mice that underwent intrahepatic injection of RIL175 cells or non-tumor vehicle controls (cecal content, 21 d-post tumor initiation). Cecal content was filtered and stored in 50% glycerol with microbiota transplantation into recipients after polyethylene glycol (PEG; laxative) treatment, with methodology based on prior transplantation studies (*57*, *82*). Fecal samples, cecal, and small intestinal (jejunal/ileal) content were collected for 16S rRNA-Seq with library preparation and sequencing completed at the NIH Laboratory of Integrative Cancer Immunology-Microbiome and Genetics Core. Samples were analyzed via QIIME2 (*83*) and LefSE (*84*) pipelines. Further details provided within Expanded Methodology.

In addition, authors performed nearest-neighbor cellular analyses from clinical HCC surgical specimens conducted at the MedStar Georgetown University Hospital (Washington DC, IRB #2017-0365). Following surgery, cellular profiling was conducted via co-detection by indexing (CODEX) imaging and accessed via the National Cancer Institute Halo Image Analysis Resource. Methodology and sample processing reported in (*36*). Pathway enrichment analyses (ReactomePA) were conducted from bulk RNA-Seq datasets and differential expressed gene lists provided in (*35*). Remaining methodology, materials, analyses, and data storage provided within Expanded Methodology.

## Supporting information

Supplemental File 1 Tumor adjacent tissue exhibits altered immune and neurological pathways

Supplemental File 2 CODEX identification markers

Supplemental File 3 scRNA-Seq differentially expressed gene list

Supplemental File 4 Neuroimmune gene list

Supplemental File 5 Microbiome Analyses

## Acknowledgments

Authors thank Sophie Wang and Gary Shaw (Greten Laboratory Managers), Vanessa Catania, Varun Subramanyam, Gauri Garg, Thien Nguyen, Aviva Leah Menashy, and Xiao Bin for procedural assistance. We thank Dr. Jay Berzofsky and colleagues from the Greten, Berzofsky, Trinchieri, and Xie laboratories for valued discussion regarding neuroimmune and microbiome interactions.

Drs. Lee Chedester (National Institute on Alcohol Abuse and Alcoholism), Luiz F. Barella (formerly National Institute of Diabetes and Digestive and Kidney Diseases), and Michael J. Reilly (Charles River Surgery Supervisor) offered technical equipment and/or guidance for hepatic vagotomy. MR3 KO breeding pairs were kindly provided by Dr. Jürgen Wess (NIDDK). Dr. David Kleiner offered pathology and CODEX interpretation guidance. Dr. William Telford at the Research Flow Facility (National Cancer Institute) reviewed immune sorting (scRNA-Seq). We thank members of the Murine Phenotyping Core (NIH-NHLBI), specifically Audrey Noguchi, Morteza Peiravi, and Heather Potts. We are grateful to Dr. Michael Kelly and members of the Single Cell Analysis Facility (NCI-SCAF), notably Jatinder Singh, Charlie Seibert, Kimia Dadkhah, and Ian Taukulis for scRNA-Seq preparation and deposition of primary data (Frederick National Lab contract #75N91019D00024). We thank Colm O’hUigin and Laboratory of Integrative Cancer Immunology-Microbiome and Genetics Core members, Wuxing (Jane) Yuan and Shah M. Rashed, for 16S rRNA library prep, sequencing, and primary data storage. Drs. Jon Inglefield and Yanyu Wang (Frederick Histopathology Core, Leidos Biomedical Research, Inc.) provided valued Luminex analyses. We thank Dr. Baktiar Karim and members of the Molecular Histopathology Laboratory at the Frederick National Laboratory including Drs. Tammy Beachley, Donna Butcher, and Tamara Morgan for cFOS staining and primary data storage. The authors thank LASP (Laboratory Animal Sciences Program) for animal support, as well as Dr. Marina Villamor-Paya for helpful discussion in mouse breeding strategy.

Data analysis utilized computation resources and/or storage of the NIH HPC Biowulf cluster (http://hpc.nih.gov), NCI Partek Flow workspace, and NIH Halo Image Analysis Resource. Graphical figures were created with Biorender (http://biorender.com/).

## Funding

KCB receives financial support through an iCURE-NCI Fellowship from the NIH (2020), Sallie Rosen Kaplan Women in Science Funding (2023), and SITC-Genentech Women in Science Fellowship (2023). B.R. is currently funded by the Deutsche Forschungsgemeinschaft (DFG, German Research Foundation) under Germany’s Excellence Strategy - EXC 2180 – 390900677. FJR-M receives funding through the NIH Medical Research Scholars Program. TFG laboratory funding provided by the Intramural Research Program of the NIH, NCI (ZIA BC011345, ZO1 BC010870). TFG and GT were support by NCI FLEX award.

## Author contributions

KCB developed project concept, optimized procedures, performed experiments, and analyzed data/prepared visualization in collaboration with members from the Greten, Trinchieri, and Xie laboratories. Manuscript was written by KCB with input from all coauthors. RT, BR, YM, M-RB, CM, MD, AN, JQ, PH, MS, BLG, SW, DAS, FJR-M, JM, SG, SNG, BD, JG, RR, and AKD designed procedures, conducted experiments and/or performed analyses. CX, GT, FK, and TFG guided laboratory members, project development, resources, and analytical interpretation. TFG led funding acquisition, collaborative supervision across research teams, conceptualized scientific hypotheses, guided research and paper development, and edited finalized manuscript.

## Competing interests

The authors declare no competing interests.

## Expanded Methodology

### Mice and vagotomy procedures

All animal procedures and experimental endpoints were approved by the NCI Institutional Animal Care and Use Committee in accordance with the National Research Council’s *Guide for the Care and Use of Laboratory Animals*. Murine procedures were listed under animal study proposal MOB-028, TGOB-013, and TGOB-015. Mice were housed at the CRC Animal Facility and NHLBI Murine Phenotyping Core on 12-hr light/dark cycles and received *ad libitum* access to food and water.

Female and male mice were purchased through Charles River Laboratories: C57BL/6 (strain code: #556), BALB/c (strain code: #028), and C57BL/6-Tg(TcraTcrb)1100Mjb/J (OT-1; strain code: #003831); or Jackson Laboratories: C57BL/6J (strain code: #000664), C57BL/6-Tg(Cd8a-cre)1Itan/J (CD8^cre^ strain code: #008766), and C57BL/6J-Rag1<em10lutzy>/J (Rag1KO strain code: #034159). MR3-flox mice were generated as described in Yamada et al., 2001 (*85*) and kindly provided by Dr. Jürgen Wess (NIH-NIDDK, Bethesda, MD 20892). To develop CD8^cre^M3R^f/f^ mice, CD8^cre^ mice were bred with MR3-flox animals and CD8^cre^MR3^WT/f^ offspring were backcrossed with MR3-flox animals. Littermate controls included CD8^cre^, M3R^f/f^, and CD8^cre^MR3^WT/f^. Breeding was performed by researchers within the CRC Animal Facility and genotyping confirmed through TransnetYX services. Procedures utilized sex-matched controls two-four months of age and hepatic or sham vagotomies performed upon 8-12-week-old mice.

Hepatic vagotomized or sham surgical controls were obtained from Charles River Laboratories (surgical services discontinued 2022). Alternatively, vagotomy was performed as described in Teratani et al., 2020 (*29*). Briefly, a vertical incision was made from the xiphoid process to mid-abdomen. The common hepatic branch was identified with surgical loupes (4.0X, 340 mm working distance) off the left (anterior) vagal branch from where it descends from the diaphragm. The hepatic branch was isolated and snipped and/or torn with surgical tweezers. For the sham surgery, the hepatic branch was identified but not cut. The abdominal cavity was flooded wither sterile 0.9% saline solution to prevent adhesions. Muscular incisions were closed with surgical sutures and skin flaps closed with wound clips with analgesic follow-up. A two-week recovery period was scheduled between surgical procedures.

### Cell lines and tumor models

Murine hepatocellular carcinoma cell line RIL175, as described in Kapanadze et al., 2013 (*86*), was cultured in RPMI 1640 media (Thermo Fisher Scientific, catalogue #61870127) +10% FCS (fecal cos serum: Corning Inc., catalogue #MT35010CV). The B16-F10 (murine melanoma) cell line was provided by Dr. Ugur Sahin (Mainz, Germany), as described in Kreiter et al., 2015 (*87*) and A20 (murine lymphoma) and B16-F10-OVA cell lines were purchased from ATCC. For in vitro studies and metastatic tumor cell lines, cell media (RPMI 1640 + 10% FCS) was further supplemented with 1 mM sodium pyruvate (Thermo Fisher Scientific, catalogue #11360070), 1mM-1M HEPES (Thermo Fisher Scientific, catalogue #15630080) +1mM NEAA (Thermo Fisher Scientific, catalogue #11140050) and/or 500 ng/mL puromycin (InvivoGen), and/or 3.5M beta-mercapthoethanol (Sigma-Aldrich catalogue #60-24-2). Penicillin-streptomycin solution 1% (Thermo Fisher Scientific, catalogue #5070063) was added to prevent bacterial contamination. Aliquots were passaged less than 12 times prior to use, see protocols listed in Ruf et al., 2023 (*36*). Cell lines were negative for contamination following routine bacterial and mycoplasma testing (May 2022, Molecular Testing of Biological Materials NCI, Frederick 21702).

Tumor cell initiation was performed as described in Chi et al., 2018 (*33*) and detailed in methodology reported by Brown et al., 2018 (*81*). For intrahepatic models, 20 µL/mouse injections comprised of 1:1 sterile PBS with 2.5×10^5^ cancer cells and Matrigel (Corning Inc., catalogue #354230) were injected into the left lateral lobe of the liver. Silver nitrate (Vet One, catalogue #NDC13895-706-10) was applied to staunch hepatic bleeding and incisions were closed with sterile sutures and wound clips. Subcutaneous (pelvis) and tail vein injections contained 100 µL/mouse of 1.0×10^6^ cancer cells in sterile PBS. Mice were randomized prior to surgical procedures. *In vivo* imaging (IVIS Spectrum®) of luciferase-expressing RIL-175 cells was performed with Xenogen IVIS 100 Imaging technology. Mice underwent imaging 7.5 minutes after intraperitoneal injection of 150 mg/kg luciferin (REGIS Technologies, catalogue #2591-17-5). Unless otherwise noted, tumor weights were measured at 21 d following initiation. Mice were only removed from analysis if the animal died prior to experimental termination (*i.e.*, during surgical procedures) or due to improper surgical initiation (*i.e.,* accidental injection into non-hepatic abdominal organs).

### Acetylcholine agonist models

Muscarinic treatment followed protocol reported in Renz et al., 2018 (*22*). Bethanechol chloride (Millipore Sigma Aldrich catalogue #1071009) was administered in drinking water (400 μg/mL). Agonist exposure began 3-5 days prior to tumor initiation and fresh water was replenished 1-2X/wk throughout experimental procedures. Mice received *ad libitum* access to drinking water. Comparable water consumption was observed in bethanechol-treated mice and water-only controls.

### CD8 T-cell depletion

Depletion of CD8+ T cells were conducted by 100 µL intraperitoneal injection containing 200 µg/mouse of antibody (InVivoMAb anti-mouse CD8α, Clone 2.43, BioxCell). Mice were injected one day prior to tumor initiation and every 7 d following until experimental termination. Isotype-matched IgG2b (InVivoMAb rat IgG2b isotype control, anti-keyhole limpet hemocyanin, Clone LTF-2, BioxCell) was administered at 200 ug/mouse to IgG controls.

### Microbiota Transplant

The cecal pouch of age-matched RIL175 tumor-bearing mice and non-tumor controls (21 d post-tumor initiation) were utilized for microbiota transplant. Individual cecal content was added into sterile 50% glycerol solutions (5 mL). After mixing, cecal content was filtered through a 100 µm filter and stored at −80°C in 1 mL aliquots. Recipient mice were treated with PEG (polyethylene glycol, Millipore Sigma catalogue #25322-68-3) to deplete intestinal microbiome utilizing a protocol for 90% reduction of cecal 16S DNA copies as reported by Wrzosek et al., 2018 (*82*). Briefly, mice received four rounds of oral gavages (200 µL with 425 g/L PEG in sterile distilled H_2_O) every 20 min. After the final PEG treatment, mice received the first CMT via oral gavage (100 µL): d 1. Oral gavage was repeated on days 3, 5, and 8. Fecal pellets were collected before and throughout experiment to ensure successful microbiota transplantation via 16SrRNA-Seq. All experiments concluded within 10 d of final oral gavage.

### Behavior and Activity Phenotyping

Mice that underwent phenotyping assays were housed in the Murine Phenotyping Core within the NHLBI (National Heart, Lung, and Blood Institute) on a 12 h light/dark cycle. All animal procedures were performed in accordance NCI Division of Intramural Research Animal Care and Use Committee (proposal TGOB-015). Prior to recorded tests, mice underwent an acclimatization and gentle handling period. Analyses were repeated at Baseline (post-vagotomy), Early (<7 d post-intrahepatic injection), and Late (> 7 d post-intrahepatic injection) for vagotomy procedures and Baseline (<7 d pre-cecal microbiota transplantation) and CMT (>7 d initial bacterial gavage).

### Phenotyper and XY Cage Monitoring

Mice underwent home cage monitoring (72 h /round) in PhenoTyper home cages (Noldus, Wageningen, The Netherlands) equipped with infrared camera top unit. Videos from the PhenoTyper cages were analyzed using EthoVision XT behavioral tracking software to measure activity and movement of the mice. Mice were singly housed during home cage monitoring and were provided with ALPHA-dri bedding and *ad libitum* access to food and water.

### Open Field Test

Mice were removed from home cages and placed individually placed into 16’ X 16’ X 16’ Perspex® arena viewing chamber for free exploration. Mice were recorded for 30 min prior to return to home cages. Distance, velocity, mobility, and time spent in defined zones were analyzed by ANY-maze behavioral tracking software (Stoelting Co, Il).

### Spontaneous Y maze

Mice were placed into a three-armed polycarbonate Y maze consisting of a start arm (18.4 cm start box connected to a 30.5 cm runway by a door, with two 36.8 cm side arms at 120° angle from each other off a central hub). Mice were placed within the start box, the door was opened and the mouse was allowed to freely explore the maze for five minutes. Total number of arm visits were scored and percent alternation (total alternations/possible triads X 100) was manually scored.

### Clinical Analyses Kaplan-Meier Plots

Kaplan-Meier plots were determined by assessing *CHRM3* on TIMER2.0 via the Kaplan-Meier plotter feature (*88*) and the Human Protein Atlas (*89*), website accessed December 2023.

### Reactome PA and RNA-Seq

To examine altered cholinergic and neurological pathways in cancer, we utilized a dataset provided in Aran et al., 2017 (*35*). Here, authors performed differential gene expression analyses on 6506 biopsies across eight tumor types assessing healthy and tumor-associated tissues, methodology provided in open access DOI: <u>10.1038/s41467-017-01027-z</u>. We performed ReactomePA analyses on published Supplementary Data 1, the differential expression analysis between healthy and tumor-adjacent (NAT) tissues with ReactomePA package (*90*), visualization created in RStudio (version 2022.07.1).

### CODEX Analyses

Co-detection by indexing (CODEX) multiplexed tissue imaging was performed on hepatocellular carcinoma (HCC) patients utilizing a 37-plex CODEX panel. Efforts were made to ensure sex and ethnic diversity during the recruitment process at MedStar Georgetown University Hospital (Washington DC). Biopsies were collected from patients undergoing surgical HCC resection. Informed consent for collection of clinical and biological data and appropriate transfer/storage of data followed Institutional Review Board protocol #2017-0365 and Material Transfer Agreement #43655-18. Raw CODEX images are hosted at The Cancer Imaging Archives (TCIA) under https://doi.org/10.7937/bh0r-y074. Clinical and pathological information, specimen preparation, antibody validation, and CODEX multiplex immunofluorescence methodology and imaging, and data location provided in Ruf et al., 2023 (*36*).

For nearest-neighbor nerve analyses, high-resolution primary images from 12 HCC specimens were reviewed from the NCI HALO Image Analysis Resource. Neuronal axons were classified as non-nucleated (DAPI-negative), non-bile duct (pan cytokeratin-negative), NCAM+ branches (neural cell adhesion molecule+, >50 µm). Three nerve-rich regions were identified in each specimen. Same-sized images (100x zoom) with distinct antibody markers were taken at each region to manually identify nearest cell neighbor. Image 1 (Structural/Stromal Panel), Image 2 (Lymphoid Panel), Image 3 (Myeloid Panel). Images (jpg, tif files) were subsequently analyzed in ImageJ (FIJI) to manually determine closest cell neighbors. A select number of protein markers was used to identify structural/stromal, myeloid and lymphoid cells (see Supplementary Table 2).

### Immune cell isolation and flow cytometry

Single-cell isolation of splenic, liver, and tumor-infiltrating cells followed reported protocols, namely Heinrich et al., 2021 (*91*) and Ruf et al., 2023 (*36*). Following euthanasia, organs were removed from animals and solid tumors (RIL175, B16-F10) were excised from the liver or processed as tumorous liver (A20). Both spleens and livers were homogenized and filtered through 70 μm nylon mesh and centrifuged at 4°C for 15 minutes (400x RCF). Livers underwent further density-gradient centrifugation with 90% Percoll (Cytiva, catalogue #17089101) to isolate immune cells from stromal components. Following homogenization, tumors were dissociated via gentleMACS Octo Dissociator (Miltenyi Biotec catalogue #130-096-427, murine tumor program) and subsequent density-gradient centrifugation with Lympholyte Cell Separation Media (Cedarlane Laboratories, catalogue #CL5035). All samples were treated with ACK-Lysis Buffer (Quality Biologicals, catalogue #118-156-721) to remove red blood cells. Single cell isolates were passed through 70 μm nylon mesh a second time to remove residual debris and samples were stored at −4°C in PBS+10% FCS prior to staining.

All staining was performed at 4°C. Murine samples were stained as follows: Live/Dead staining using Zombie UV(TM) Fixable Viability (BioLegend, catalogue #423108) for 20 min in PBS and subsequent Fc-blocking (eBiosciences, catalogue #14-0161-85) for 10 min in FACS buffer. FACS buffer was comprised of 9.7%10X PBS (Corning Inc, catalogue #46-013-CM), 20% FCS, and 0.1% sodium azide (Sigma-Aldrich, catalogue #26628-22-8). Surface staining was conducted for 30 min, antibodies include: anti-CD45 on Alexa Fluor 700 (BioLegend, catalogue #103128, clone #30-F11), anti-Ly6G on Alexa Fluor 700 (BD Biosciences, catalogue #561236, Clone #1A8), anti-CD4 on BV605 (BioLegend, catalogue #100547, clone #RM4-5), anti-Ly6C on APC/Cy7 (BioLegend, catalogue #128026, clone #HK1.4), anti-CD3 on PE (BioLegend, catalogue #100206, clone#17A2), anti-CD19 on PerCp-Cy5.5 (BioLegend, catalogue #152406, clone #1D3/CD19), anti-F4/80 on FITC (BioLegend, catalogue #123108, clone #BM8), anti-CD11b on Pacific blue (Biolegend, catalogue #101224, clone#M1/70), anti-TCRβ on PE/Cy7 (BioLegend, catalogue #109222, clone #H57-597), anti-MHCII on BV510 (BioLegend, catalogue #107635, clone#M5/114.15.2), anti-CD11c on BV650 (BioLegend, catalogue #117339, clone #N418), anti-CD8 on BV785 (BioLegend, catalogue #100749, clone #53-6.7), anti-CD45R/B220 on Alexa Fluor 700 (BioLegend catalogue #103232, clone #RA3-6B2), anti-F4/80 on Alexa Fluor (BioLegend catalogue # 123130, clone #BM8), anti CD11b on Alexa Fluor (BioLegend catalogue #101222, clone #M1/70), anti-CD3 on Alexa Fluor 594 (Biolegend catalogue #100240, clone #17A2), anti-TCRβ on APC/Fire^TM^ 750 (Biolegend, catalogue #109246, clone #H57-597), anti-NK1.1 (BD Biosciences, catalogue #741926, clone #PK136 [RUO]), and anti-PD1 on FITC (BioLegend, catalogue #135214, clone#29F.1A12), and Cd1d tetramer on APC (NIH tetramer core facility in collaboration with Emory University).

Prior to intracellular cytokine staining, cells underwent PMA/ionomycin stimulation. To stimulate we utilized Leukocyte Activation Cocktail, with BD GolgiPlug™ (BD Biosciences catalogue #550583) comprised of PMA/ionomycin and Brefeldin A. More specifically, cells underwent 4 h stimulation with media containing 0.4 µg/mL IL2 and/or Leukocyte Activation Cocktail, with BD GolgiPlug™ (20 µL/mL media). Prior to intracellular staining, cells were fixed using the BD Cytofix/Cytoperm™ Fixation/Permeabilization Kit (BD Biosciences, catalogue #554714) or the BD Pharmingen™ Transcription Factor Buffer Set (BD Biosciences, catalogue #562574 [for FoxP3 staining]) following manufacturer’s recommendation. Subsequent intracellular cytokine staining (30 min) utilized the following antibodies: anti-perforin on APC (BioLegend, catalogue #154304, clone#S16009A), anti-IL6 on APC (BioLegend, catalogue #504508, clone#MP5-20F3), anti-TNFα on PerCP-Cy5.5 (BioLegend, catalogue #506322, clone#MP6-XT22), anti-granzyme B on FITC (BioLegend, catalogue #515403, clone#GB11), anti-IFNγ onPE/Cy7 (BioLegend, catalogue # 505826, clone#XMG1.2), anti-IL10 on BV421 (Biolegend, catalogue #505022, clone#JES5-16E3), Upon staining, samples were assessed on a CytoFLEX LX flow cytometer (Beckman Coulter, RRID:SCR_0129627). Subsequent analyses were performed utilizing FlowJo software (vs 10, RRID:SCR_008520) or Partek Flow (NIH HPC Biowulf). Frequencies (% Live cells) or cell counts (absolute cells: population frequency x total live cells / sample weight) provided. The absolute number of immune cells was calculated by multiplying frequency by the total live cells and then divided by liver weight.

### scRNA-Seq

#### Cell sorting, library preparation, and 10X genomics sequencing

Eight samples were prepared for scRNA-Seq. Solid RIL175 tumors were excised from SV and HV livers (n = 3 / group) 21 d following tumor initiation. RIL175 tumor-bearing mice (n = 3 SV, 3 HV mice) from non-tumorous liver and matched (pooled) tumors. Immune cells from tumors and livers were isolated as described in earlier methodology. Following surface staining, Live+CD45+ cells were sorted into PBS + 0.04% BSA (ThermoFisher Scientific, catalogue #B14) at the NIH Flow Cytometry Core using the Aurora CS Cell Sorter (Cytek® Biosciences, RRID:SCR_XXXXX). Subsequent cell counts and viability were determined with propidium iodide/acridine staining using the Luna FL Automated Fluorescence Cell Counter (Logos Biosystem).

Samples were provided to SCAF (NIH Single Cell Analysis Facility) for library preparation, sequencing, and data processing. Cell counts were determined via LunaFx7 fluorescent cell counter (Logos Biosystem) and adjusted cell counts targeting 6,000 cells/lane were prepared. Library preparation and sample loading were performed by SCAF and in accordance with the 10X Genomics Single Cell User Guide utilizing Chromium Connect (10X Genomics automation platform).

Sequencing was performed using the NovaSeq 6000 Reagent Kit (Illumina). Two sequencing runs (NovaSeq 6k S1 100-cycle) were performed via 5’ Immune Profiling v2 Dual-Index chemistry (10x Genomics).

#### scRNA-Seq Processing

Base calling of data and quality scoring utilized RTA 3.4.4 (Illumina). Subsequent data processing was performed with Cell Ranger v7.1.0 (Illumina), including demultiplexing (Bcl2fastq 2.20.0) and alignment (STAR 2.7.2a) onto mouse genomics reference (refdata-gex-mm10-2020-A). Following processing, Mean Reads per Cell = 83, 551, Total Genes Detected = 19,745, and Median UMI Counters per Cell = 6,237 (values averaged across all sample outputs). The sequence saturation ranged from 81%-93% indicating a robust mapping of sequence data to the mouse transcriptome.

#### scRNA-Seq Analyses

Data was placed onto Biowulf (NIH HPC Linux cluster) and downstream analyses was performed using Seurat package (vs 4.3.0.1) under RStudio (vs 2023.09.1+494). The count matrix files were read using Read10X_h5 function, then converted into Seurat objects excluding genes expressed less than 10 cells. Low quality cells were removed by the following criteria: cells with features fewer than 500 or above 4000 genes; nCounts fewer than 500 or above 14000; log_10_(features/counts) <0.8; mitochondrial RNA content above 20%. Doublets in each sample were removed using DoubletFinder (vs 2.0.3). Normalization was performed using SCTransform function in individual samples, then all samples were merged into a single Seurat object. Harmony (vs 0.1.1) was applied for batch correction.

The FindClusters function was used to find appropriate clusters at resolution 0.4 to create separate clusters of T, B, NK and Myeloid cells. Further subclustering of T cells was performed using the FindClusters function at resolution 0.45. Differential gene analysis was completed for both all clusters, using FindAllMarkers(), as well as to separate specific memory clusters, using FindMarkers(), that persisted across multiple resolutions as independent at thresholds of min.pct at 0.1,logfc.threshold at 0.1, only.pos as TRUE. UMAPs were created using the RunUMAP function. Differential gene expression was plotted using the Enhanced Volcano package (version 1.20.0)

#### AChR+ and AChR-determination

scGSEA was completed for AchR determination based on expression of all Acetylcholine Receptor genes (*Chrm1*, *Chrm2*, *Chrm3*, *Chrm4*, *Chrm5*, *Chrna1*, *Chrna2*, *Chrna3*, *Chrna4*, *Chrna5*, *Chrna6*, *Chrna7*, *Chrna8*, *Chrna9*, *Chrna*, *Chrnb1*, *Chrnb2*, *Chrnb3*, *Chrnb4*, *Chrnd*, *Chrne*, *Chrng*). Based on resulting violin plot of gene expression with bimodal values, an appropriate threshold was chosen of 0.05 as shown.

#### Pathway Enrichment Analyses

The clusterProfiler (version 4.10.0) package was used for pathway enrichment analysis with a mouse database (org.Mm.eg.db) for liver and tumor, separately. Cutoffs for clusterProfiler were p value of 0.05, function enrichGO, biological processes (BP) were chosen.

#### Monocle Trajectory Analyses

Monocole and Pseudotime analysis were completed using the Monocle3 package (version 1.3.4). For Monocle, Leiden clustering method was chosen at resolution 7e-4.

### In vitro ACh signaling assays

#### Intracellular activation assay

Lymphocytes were expanded from mouse splenocytes treated with ACK Lysing buffer. CD3+ T cells were expanded via 1 μg/ml anti-mouse CD3 (eBiosciences, catalogue #16-0031-82, clone #145-2C11) and 0.1 μg/ml anti-mouse CD28 (eBiosciences, catalogue #16-0281-85, clone #37.51) antibodies and activated with IL2 (100 ng/mL) or 3:1 ratio of CD3/CD28 Dynabeads (ThermoFischer Scientific, catalogue #11452D) to CD3+ T cells. Following 48-h expansion, lymphocytes underwent pharmaceutical exposure (2 h) followed by activation with pharmaceutical exposure (4 h). The 4-h activation consisted of 0.4 µg/mL IL2 and/or Leukocyte Activation Cocktail, with BD GolgiPlug™ (20 µL/mL media) as reported above. Pharmacological compounds included ACh chloride (Sigma Aldrich, catalogue #A2661-25G), bethanechol chloride (Sigma Aldrich, catalogue #071009-200MG), or 4-DAMP (Sigma Aldrich, catalogue #SML0255-50MG) in a dose-dependent manner. Pharmacological concentrations determined from tissue ELISA and published mouse (*22*) and *in vitro* analyses (*92*). Intracellular cytokine and cytotoxic expression levels were assessed via flow cytometry.

#### Cytotoxicity assay

B16-F10-OVA cells were labelled with CellTrace™ Violet Cell Proliferation Kit (Invitrogen, catalogue #C34557) according to manufacturer specification and plated at 1.5×10^4^ cells/well within a 96-well plate. After 12-h incubation, CD8+ T cells were cocultured at reported effector-to-target ratios. These CD8+ T cells were expanded from OT-1 splenocytes for five days with or without ACh (1 µg/ml) and enriched for CD8+ T cells with the CD8a+ T Cell Isolation Kit, mouse (Miltenyi Biotec, catalogue #130-104-075) according to manufacturer protocol. Cocultures were plated with or without ACh (1 µg/ml) for 2 d. Following coculture, tumor cells were detached and combined with collected supernatant. Live/dead staining with Zombie UV™ Fixable Viability Kit was performed to assess viability. Heat-killed cocultures provided controls for maximum cytotoxicity. Percentage of maximum cytotoxicity was calculated using the formula [(number of live cells in the control – number of live cells in the experimental condition)/(number of live cells in the control – number of live cells in the heat-killed control)].

In separate, equivalent cocultures cells were treated with Brefeldin A (Biolegend, catalogue #420601) for 4 h, following the instructions provided by the manufacturer. Intracellular cytokine and effector molecule expression of CD8+ T cells was assessed via flow cytometry.

#### MTT assay

Tumor cell lines were plated at 10,000 cells / well in a 96 well plate in tumor cell line media (RPMI +10% FCS +1% penicillin-streptomycin solution) with indicated concentrations of ACh or bethanechol, as well as media controls. After 48 h, media was removed and viability determined via an MTT Cell Proliferation Kit (Abcam, catalogue # ab211091) following manufacturer’s guidelines.

#### ELISA

Acetylcholine content was measured via the QuickDetect™ ACh (Mouse) ELISA Kit (Biovision, catalogue #E4453). Murine tissues were collected and placed into 2 mL Lysing Matrix D tubes (MP Biomedicals, catalogue #116913050-CF) containing 1 mL PBS and Roche cOmplete™ EDTA-free Protease Inhibitor (Millipore Sigma, catalogue # 11873580001). Tissues were homogenized with FastPrep®-24 (MP Biomedicals) 2 rounds (5.5 m/s, 45 sec). Homogenate was centrifuged at 4°C and chilled supernatant was collected and stored at −80°C for analyses. Data was normalized to tissue weight or protein content, as measured via Pierce™ BCA Protein Assay Kits (ThermoFischer Scientific, catalogue #23225).

#### RNA isolation and RT-qPCR

Colonic tissue (10 cm proximal colon) was excised and tissue RNA was isolated using RNeasy Mini Kit (Qiagen, catalogue #74106). RNA content and purity was determined with nanodrop (ThermoFisher Scientific, catalogue #ND-2000). Adjusted RNA content was utilized for cDNA preparation using iScript cDNA Synthesis Kit (Bio-Rad, Cat #1708891) via PCR (Bio-Rad, catalogue #1852148). Samples were stored at −80°C prior to RT-qPCR. Samples were prepared using the SsoAdvanced Universal SYBR Green Supermix (Bio-Rad, Cat #1725271) prior to RT-qPCR. All primers listed in Brown et al., 2015 (*93*). *Gadph* provided an endogenous housekeeping control and was used for normalization of ddCT calculations.

#### cFOS Staining

Mice were euthanized by displacement of air with 100% carbon dioxide and within 5 minutes decapitated for tissue collection. Brain was collected at the level of brain stem containing AP region. Brain was fixed in 10% neutral buffered formalin for a minimum of five days. After fixation, brain tissues were processed on the Sakura® Tissue-Tek® VIP™ automatic processor and embedded into paraffin blocks using Sakura® Tissue-Tek® TEC™ embedding center.

All blocks were sectioned on a manual MIRCROM HM 325 microtome and placed on charged slides. Slides were dried in an 80°C oven for one hour prior to H&E staining. Hematoxylin and Eosin (H&E) staining was performed using the Sakura® Tissue-Tek®Prisma™ automated Stainer. The slides were hydrated and stained with commercial hematoxylin, clarifier, bluing reagent and eosin-Y. A regressive staining method was used. This method intentionally overstains tissues and then uses a differentiation step (clarifier/bluing reagents) to remove excess stain. The slides were cover slipped using the Sakura® Tissue-Tek® Glas™ automatic cover slipper and dried prior to review. H&E slides were evaluated by one pathologist to confirm the presence of AP region. IHC staining was performed on 5 μm tissue section using LeicaBiosystems’ BondRX autostainer with the following conditions: Epitope Retrieval 1 (Citrate) 20’, Normal Goat Serum Block, c-Fos (Synaptic Systems #226 308, 1:1000 incubated 30’), Biotinylated Goat anti-Guinea Pig IgG (Vector Labs), Streptavidin-HRP (Invitrogen #434323), and the Bond Polymer Refine Detection Kit (LeicaBiosystems #DS9800) minus the PostPrimary and Polymer Reagents. Buffer was used in place of the primary antibody for the negative control. Slides were removed from the Bond autostainer, dehydrated through graded ethanols, cleared with xylenes, and coverslipped.

#### 16SrRNA Sequencing

The following samples were utilized for 16SrRNA-Seq: fecal pellets, small intestine (ileal/jejunal content collected 10 cm of the distal small intestine), cecum (cecal content), and donor content (frozen bacterial slurries in 50% glycerol). Prior to libarary preparation, samples were added into pre-heated (10 min, 60°C) MBL Solution (Qiagen) according to the NIH Laboratory of Integrative Cancer Immunology-Microbiome and Genetics Core (LICI-MGC) pipeline.

Library preparation and sequencing was performed by the LICI-MGC Core following established protocols. DNA extraction utilized the PowerSoil DNA Isolation Kit (Qiagen, catalogue #47014). Bacterial amplification of the 16S V4 region (515F: 5′-GTGCCAGCAGCCGCGGTAA-3′, 806R: 5′-GGACTACCAGGGTATCTAAT-3′) was performed on epMotion 5073 (Qiagen) robots. 16S DNA was sequenced on the NGS MiSeq platform (Illumina) producing paired-end reads (2×250 base pairs).

Subsequent demultiplexing, quality control filtering, alpha rarefaction, and phylogenetic diversity analyses were performed with QIIME2 vs. 2023.7 (CITE). Forward reads were brought into QIIME2, the DAD2 algorithm (CITE) provided quality control and feature table construction. Sampling depth was determined based on alpha rarefaction and QIIME2 table.qzv outputs. Taxonomic classification utilized a Naïve Bayes classifier trained on dataset-specific reference reads and SILVA 132 (99% OUT sequence identity) reference taxonomy (CITE!). Linear discriminant analysis Effect Size (LefSe) analyses and visualization was conducted on the Huttenhower Galaxy server galaxy.biobakery.org). Additional data visualization was performed in R (vs. 4.2.1) with RStudio (vs. 2022.07.1) and GraphPad Prism (vs. 9+).

Raw 16SrRNA-Seq .fasq files were deposited into NCBI Sequence Read Archive (SRA) public database PRJNA105510.

#### Statistical Analyses

Sample sizes for experimental procedures were guided by previous studies and prior analyses utilizing identical tumor models (*34*, *36*, *91*). Statistical tests and outcomes are reported in the manuscript and figure legends. Analyses were conducted using GraphPad Prism vs 9+ (RRID:SCR_002798) and R (vs. 4.2.1; RStudio vs. 2022.070.01), with statistical significance set at *P* <0.05.

#### Data Availability

At this time, data requests are to be directed to Dr. Tim F. Greten.

---

**Figure S1.**
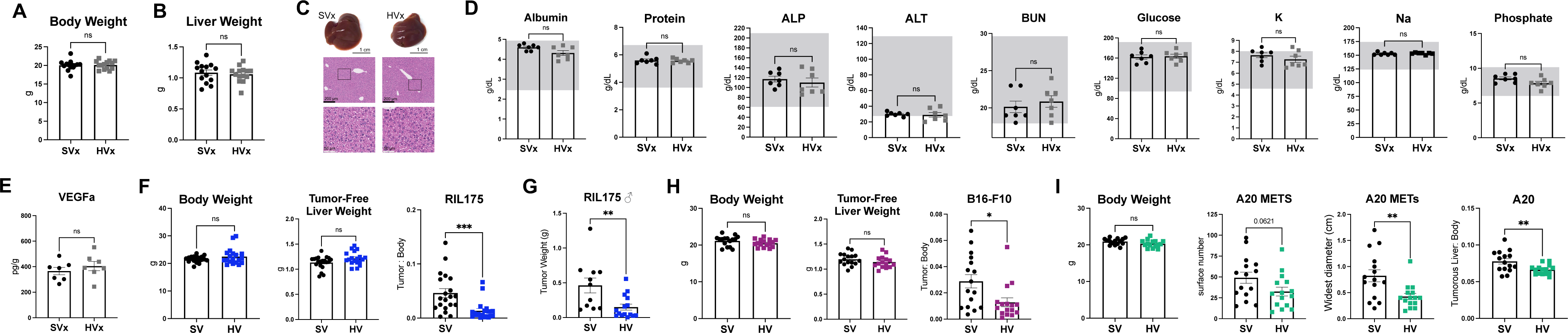
Hepatic vagotomy influences liver tumors in multiple mouse models independent of metabolic alteration. (**A**) Body weights from non-tumor bearing sham (SVx) and hepatic vagotomy (HVx) mice, *n* = 14 SVx, 14 HVx. (**B**) Liver weights from SVx and HVx mice. (**C**) Gross and H&E histology of representative SVx and HVx livers. (**D**) Liver functional assay from mouse sera in non-tumor bearing mice (*n* = 7 SVx, 7 HVx). Gray overlay indicates the reference range. (**E**) VEGFa cytokine levels from RIL175 HVs and SVs liver tissue, data normalized to tissue weight (*n* = 7 SVx, 7 HVx). Mice in **C**-**E** from cohort reported in **A** and **B**. (**F**) (*Left-right*) mouse body weight, tumor-free liver weight, and tumor:body ratio in RIL175 model (*n* = 20 SV, 19 HV). (**G**) Tumor weight in male RIL175 mice, 21 days after tumor initiation (*n* = 11 SV, 15 HV). (**H**) (*Left-right*) mouse body weight, tumor-free liver weight, and tumor:body ratio in B16-F10 model (*n* = 16 SV, 15 HV). (**I**) (*Left-right*) mouse body weight, total number of surface metastatic tumors (METs) in A20 model, diameter of largest A20 MET measured at the widest region, and A20 tumorous liver:body weight. METs measurements determined by a blinded researcher (*n* = 15 SV, 15 HV). (**I**) Data in **A**, **B**, and **F**-**I** comprised of two independent pooled experiments. Bar graphs represent mean ± SEM. Statistical significance determined by unpaired Student’s *t* test: *P < 0.05; **P < 0.01; ***P < 0.005; ns = not significant.

**Figure S2.**
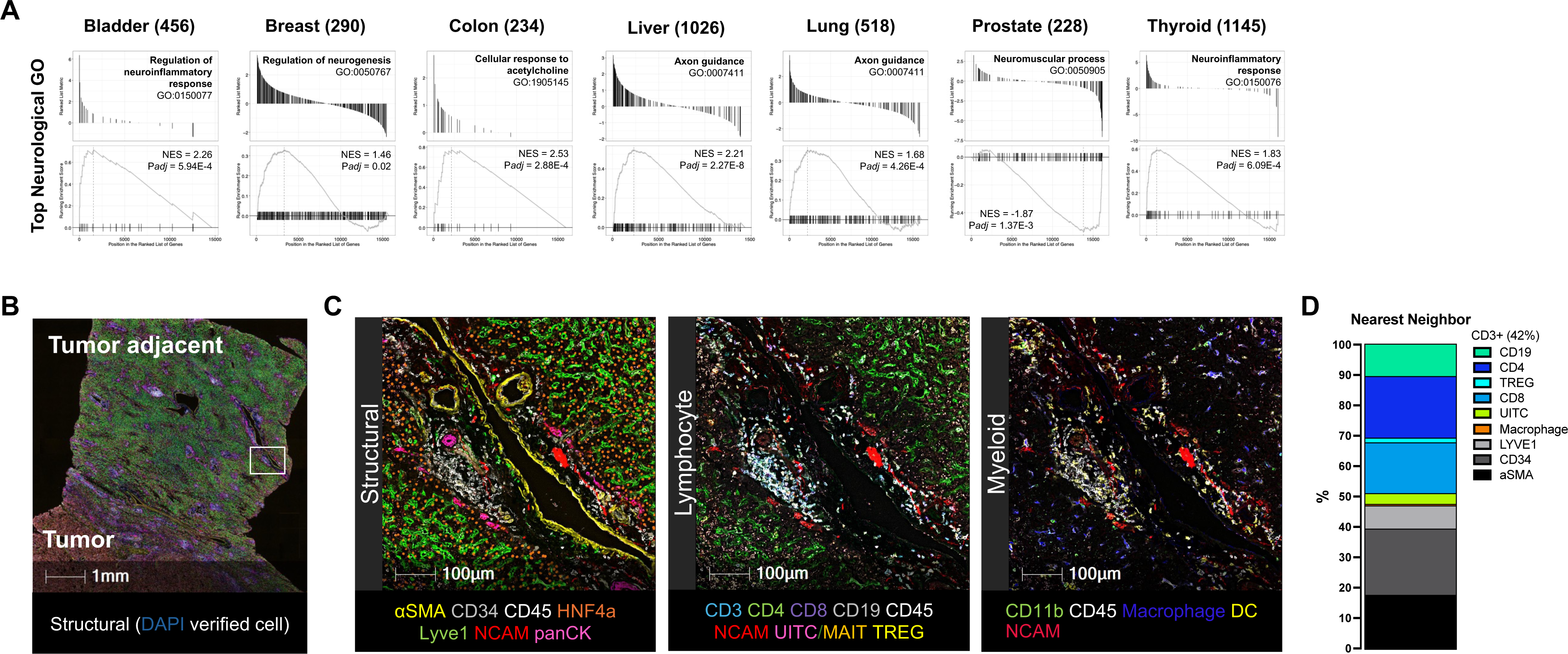
Neuroimmune interactions present across cancer populations. (**A**) Pathway enrichment analyses across seven clinical cancers from healthy tissue and tumor-adjacent specimens, analyzed from differentially expressed gene (DEG) analyses from TCGA and GTex datasets, reported in (*35*). Top neurological GO pathway from analyses as ranked by *Padj* value. Cancer types listed across the top with the total number of differentially expressed pathways within parentheses, see Supplemental File 1 for complete pathway enrichment analyses. (**B**) Example of non-magnified CODEX graph, white square indicates subsequent magnification (100X, see **fig. S2C**). (**C**) Representative **structural** (HNF4a, LYVE1, NCAM, pan cytokeratin, αSMA, CD34, CD45), **lymphocyte** (NCAM, CD45, CD3, CD19, CD4, FoxP3, CD25, CD161, TCRVα), and **myeloid** (NCAM, CD45, CD11b, CD68, CD11c) staining from biopsy reported in B. Visualized color and cell identity listed in Supplemental File 2. Cell nucleus identified by DAPI staining. (**D**) Average nearest neighbor across 12 HCC biopsy samples as determined by CODEX imaging. CODEX datasets and antibody clones provided in (*36*). αSMA = α-smooth muscle actin, EpCAM = epithelial cellular adhesion molecule, GTex = Genotype-Tissue Expression, LYVE1 = lymphatic vessel endothelial hyaluronan receptor 1, NCAM = neural cell adhesion molecule, panCK = pan-Cytokeratin, TCGA = The Cancer Genome Atlas, TREG = regulatory T cell, and UITC = unconventional and innate-like T cells (included MAIT = mucosal-associated invariant T cells).

**Figure S3.**
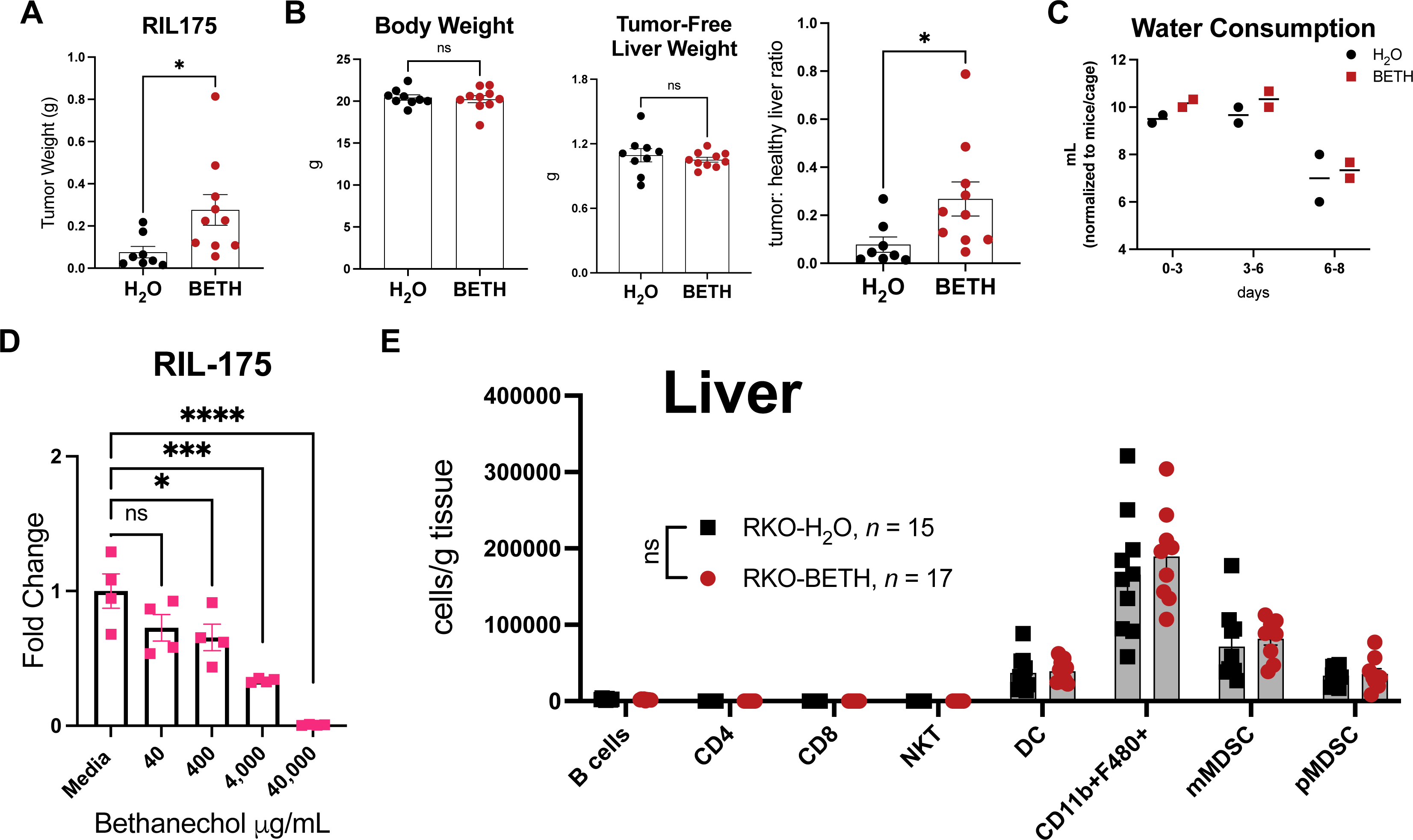
Bethanechol treatment alters liver tumor growth. (**A**) RIL175 tumor weights in H_2_O and BETH mice 21 d following intrahepatic injection. (**B**) (*Left-right*) Body weight, tumor-free liver weight, and tumor:healthy liver ratio from mouse cohort in **A**. Data pooled from two independent experiments, *n* = 8 H_2_O, 10 BETH. (**C**) Water consumption at three different timepoints in H_2_O or BETH (bethanechol: 400 μg/mL in drinking water) cages. Each dot represents a mouse cage, consumption normalized to mice per cage. (**D**) Results from colorimetric MTT assay following 48 h bethanechol treatment on RIL175 cells, four technical well replicates. Data normalized to media well controls. (**E**) Data from an independent BETH study in Rag1KO mice flow cytometry assessing myeloid immune profiles in liver. Cell counts normalized to tissue weight. Statistical significance determined by unpaired Student’s *t* test, or one-way ANOVA with *posthoc* Dunnett’s test (**D**). Bar graphs represent mean or mean ± SEM: \**P* < 0.05; \*\*\**P* < 0.005; \*\*\*\**P* < 0.001, ns = not significant.

**Figure S4.**
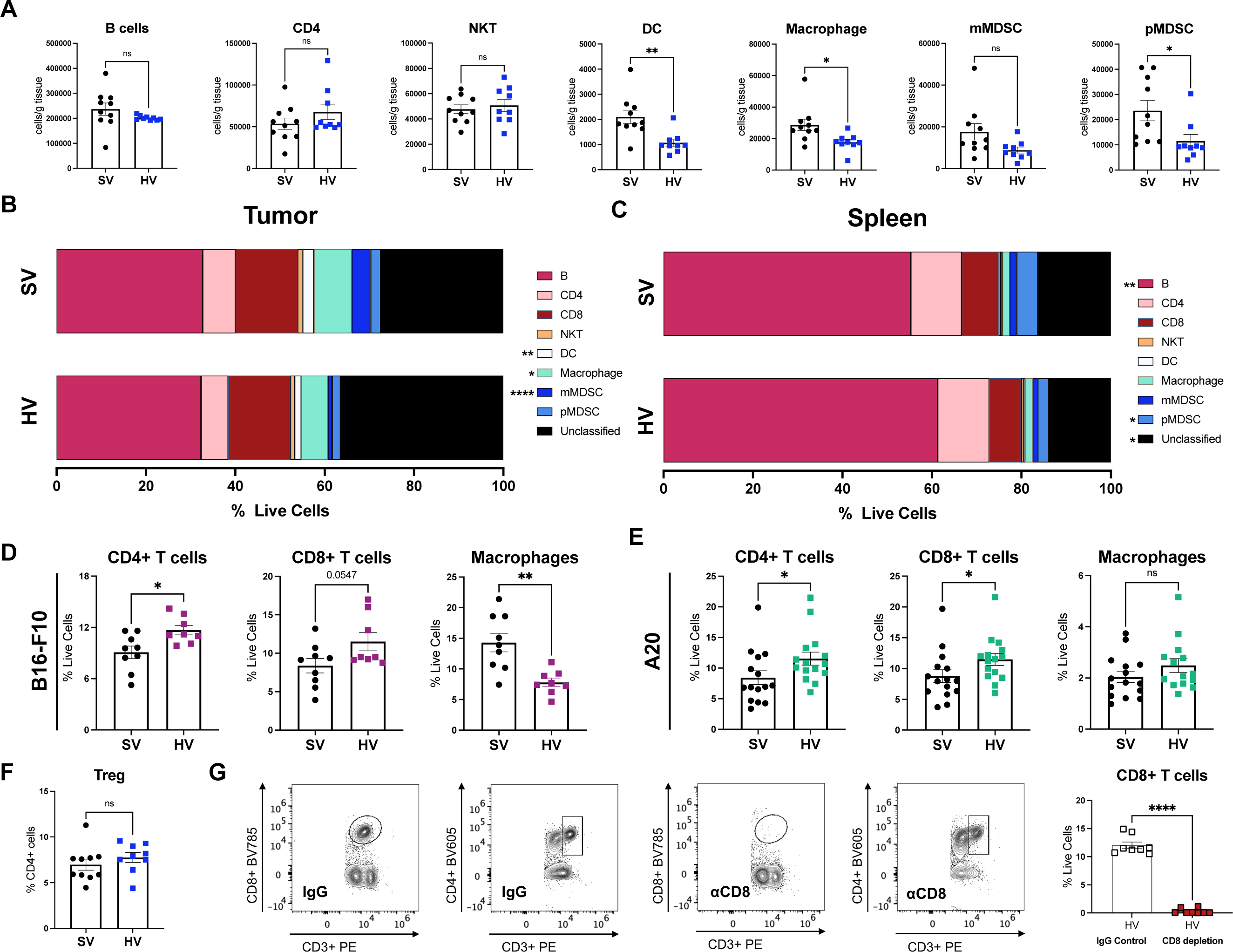
Hepatic vagotomy alters liver immune profiles. (**A**) Hepatic immune profiles assessed via flow cytometry, cell counts normalized to tissue weight at 21 d following RIL175 tumor initiation, *n* = 10 SV, 9 HV. (**B**) Average frequency (% live cells) of tumor and (**C**) splenic immune cell composition assessed via flow cytometry. (**D**) (*Left-right*) Frequency of Live+ cells of CD4+ T cells (CD3+CD4+), CD8+ T cells (CD3+CD8+), and macrophages (CD3-CD11b+F480+), 21 d following initiation of B16-F10 (*n* = 9 SV, 8 HV) or (**E**) A20 model (*n* = 15 SV, 15 HV). Mice in **E** from two independent cohorts. (**F**) Average frequency of Treg cells (CD3+CD4+FoxP3+) in RIL175-bearing SV and HV livers. Mice in **A**-**C** and **F** from the same cohort. (**G**) Anti-CD8+ T cell depletion was verified by flow cytometry, representative flow plots reported and frequency (% live cells) in HV livers reported. Bar graphs represent mean ± SEM. Statistical significance determined by unpaired Student’s *t* test: reported value 0.05<*P*<0.1, \**P* < 0.05; \*\**P* < 0.01; \*\*\*\**P* < 0.01; ns = not significant.

**Figure S5.**
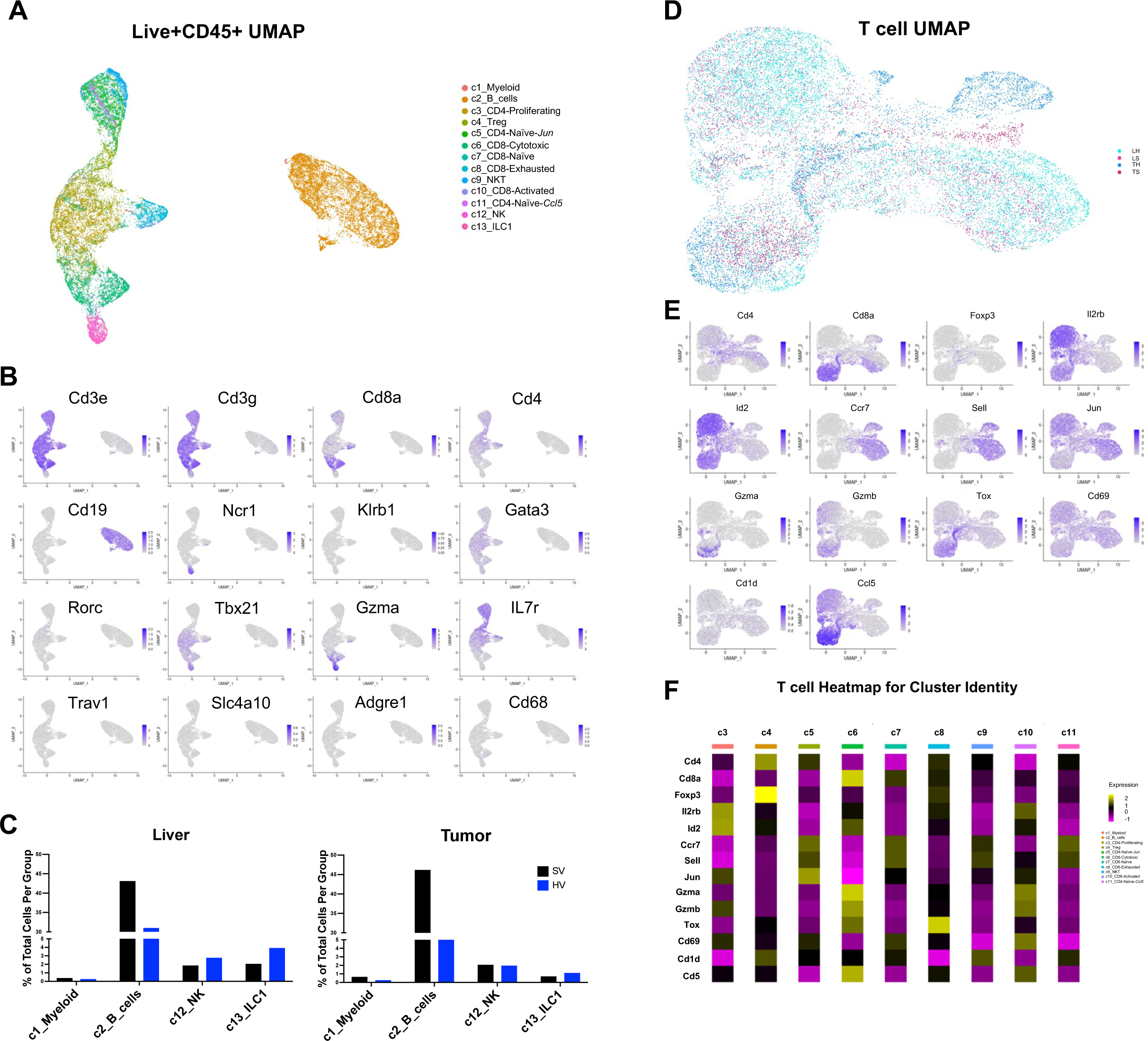
scRNA-Seq reveals distinct HV lymphocyte profiles. (**A**) UMAP and immune clusters of all liver (*n* = 3 / group) and matched, pooled RIL175 tumor samples, data from SV and HV immune (sorted Live+CD45+) cells see also **Supplemental File 3**. (**B**) Expression of critical immune markers utilized to identify subclusters within the immune UMAP, see also **Supplemental File 3**. (**C**) Percent of total cells within myeloid, B, NK, or ILC1 cell clusters from liver (tumor adjacent) and tumor (tumor-infiltrating) samples. (**D**) T cell UMAP (Live+CD3+ subsets) identified by tissue (L = liver, T = tumor) and vagotomy status (H = hepatic vagotomy, S = sham vagotomy). (**E**) Expression and (**F**) heatmap of critical immune markers utilized to identify subclusters within the T cell UMAP see also **Supplemental File 3**.

**Figure S6.**
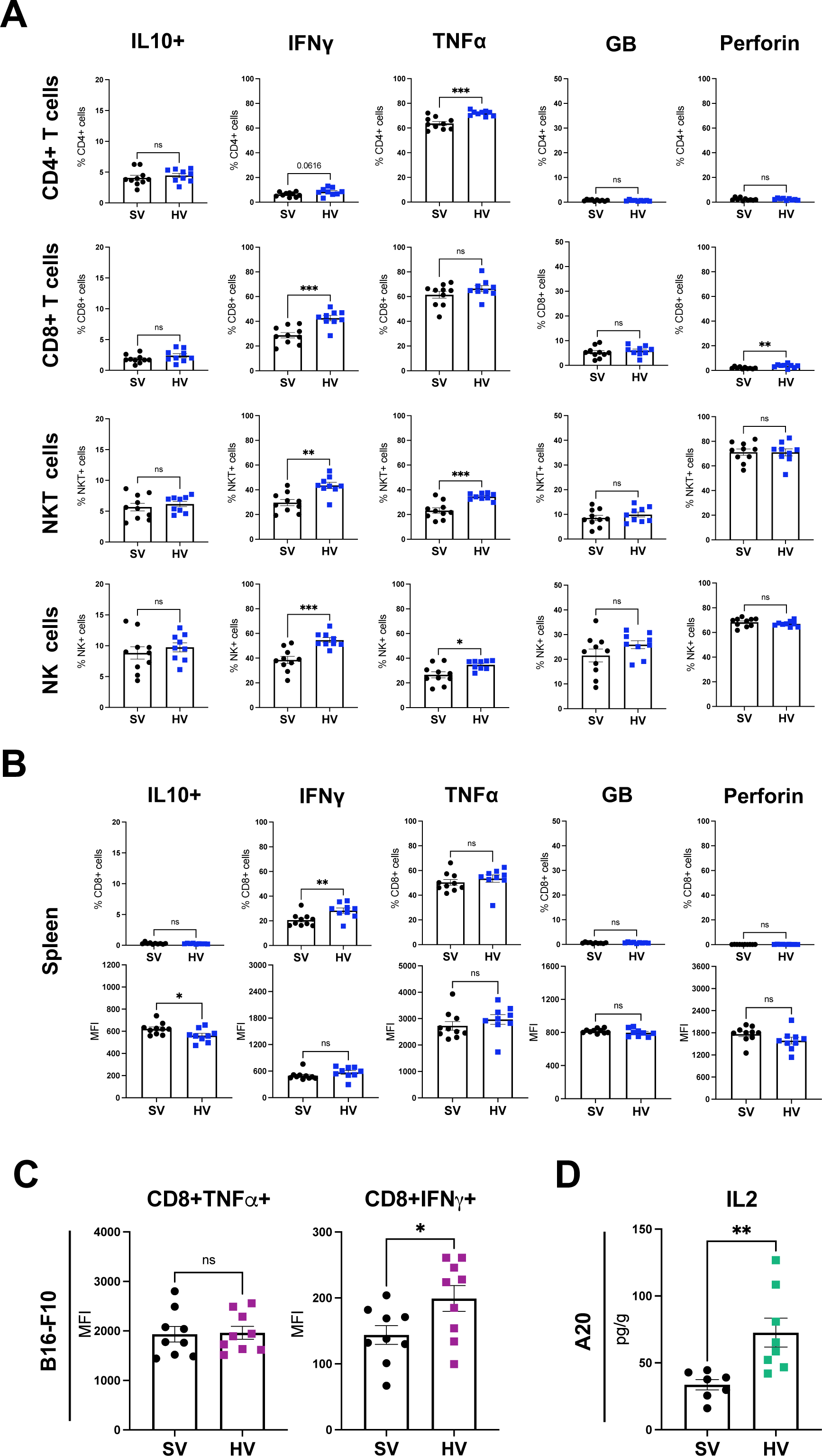
T cell stimulation reveals distinct HV lymphocyte responses. (**A**) Parental frequency of cytokine and cytotoxic markers in *ex vivo* hepatic lymphocytes as measured by flow cytometry. Samples collected 21 d following RIL175 tumor initiation and stimulated for 4 h under PMA/ionomycin (*n* = 10 SV, 9 HV). (**B**) Frequency (*top*) and MFI (*bottom*) of cytokine and cytotoxic markers from matched splenic CD8+ T cell samples. (**C**) MFI reported following intracellular cytokine staining (IFNγ, TNFα) of CD8+ T cells in B16-F10 model. Stimulation and staining as reported in **A** and **B**. (**D**) IL2 cytokine analysis from hepatic A20-tumor tissue, data normalized to hepatic tissue weight (*n* = 7 SV, 8 HV). Cytokine levels in **D** determined via a customized Procartaplex Luminex immunoassay. Bar graphs represent mean ± SEM. Statistical significance determined by unpaired Student’s *t* test: 0.05<*P*<0.1; \**P* < 0.05; \*\**P* < 0.01; \*\*\**P* < 0.005; ns = not significant.

**Figure S7.**
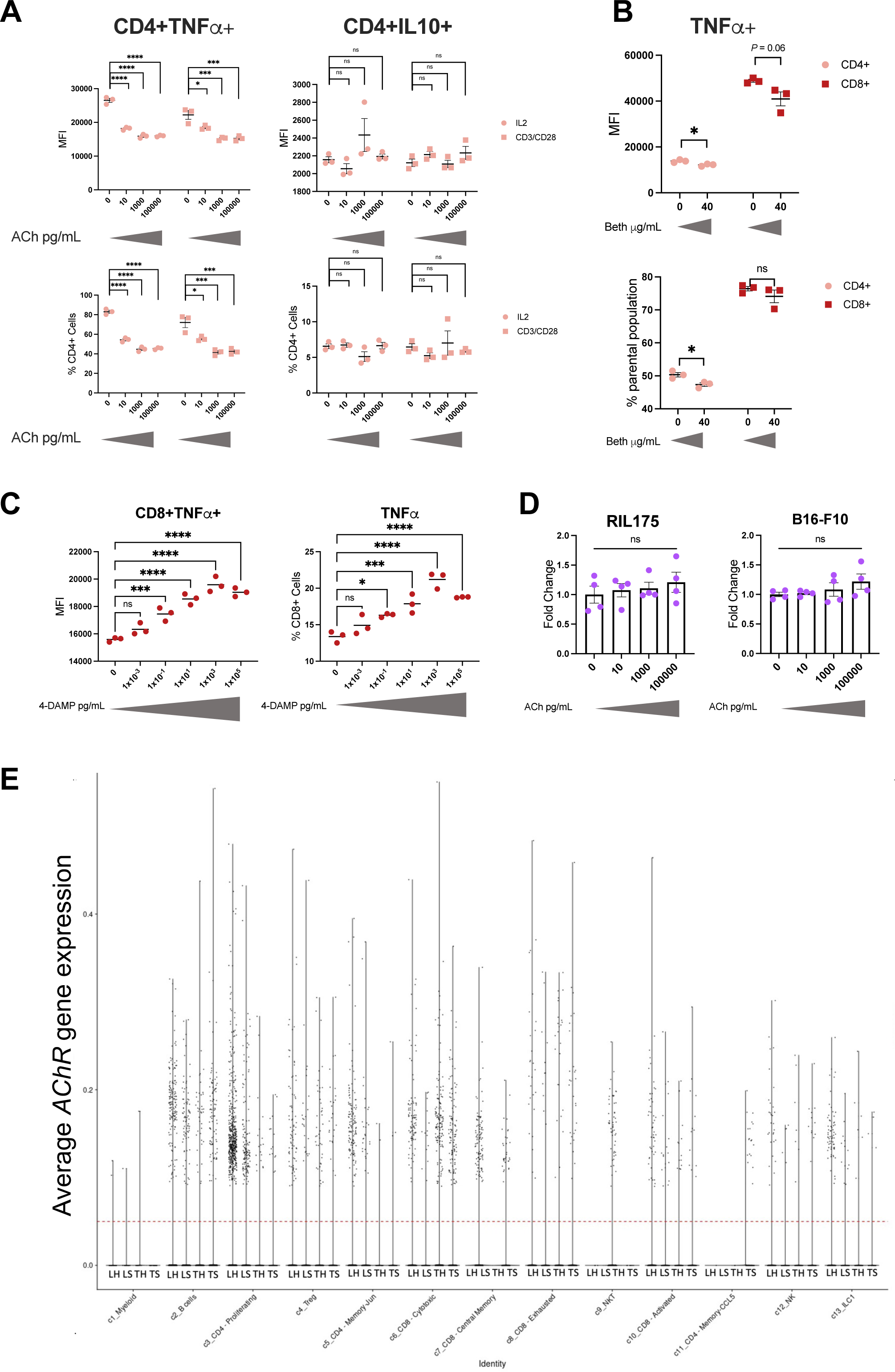
Cholinergic signaling suppresses T cell inflammation. (**A**) Splenic CD4+ T cell populations after expansion under IL2 (100 ng/mL) or CD3/CD28 Dynabeads (3:1 T cells: Dynabeads) stimulation for 48 h. TNFα+ and IL10+ intracellular cytokine staining following PMA/ionomycin activation in CD4+ T cells exposed to increasing ACh concentrations, MFI (*above*) and parental frequency (*below*) reported, *n* = 3 technical replicates. Further stimulation procedure reported in Expanded Methodology. (**B**) TNFα+ intracellular staining in lymphocytes exposed to bethanechol or (**C**) 4-DAMP (1,1-dimethyl-4-diphenylacetoxypiperidinium iodide: AChR M3 antagonist). Stimulation procedures following protocol in **A**, *n* = 3 technical replicates. (**D**) MTT assay results following 48 h ACh treatment of RIL175 and B16-F10 tumor cell lines, *n* = 4 technical replicates. The MTT assay provides a proxy of cellular viability and proliferation. Data normalized to media well controls. (**E**) Violin plot of *AChR+* and *AChR-* subsets determined by expression of muscarinic and nicotinic AChR genes. *AChR+* cells (above dotted line) identified as combined *AChR* expression > 0.05, clusters divided by tissue (L = liver, T = tumor) and vagotomy status (H = hepatic vagotomy, S = sham vagotomy), *n* = 3 / group (liver) or matched, pooled (tumor). Statistical significance determined by unpaired Student’s *t* test (**B**) or one-way ANOVA with *posthoc* Dunnett’s test (**A**, **C**, and **D**). Data in non-violin plots indicate mean and presented whiskers represent mean ± SEM: \**P* < 0.05; \*\**P* < 0.01; \*\*\**P* < 0.005; \*\*\*\**P* < 0.001, ns = not significant.

**Figure S8:**
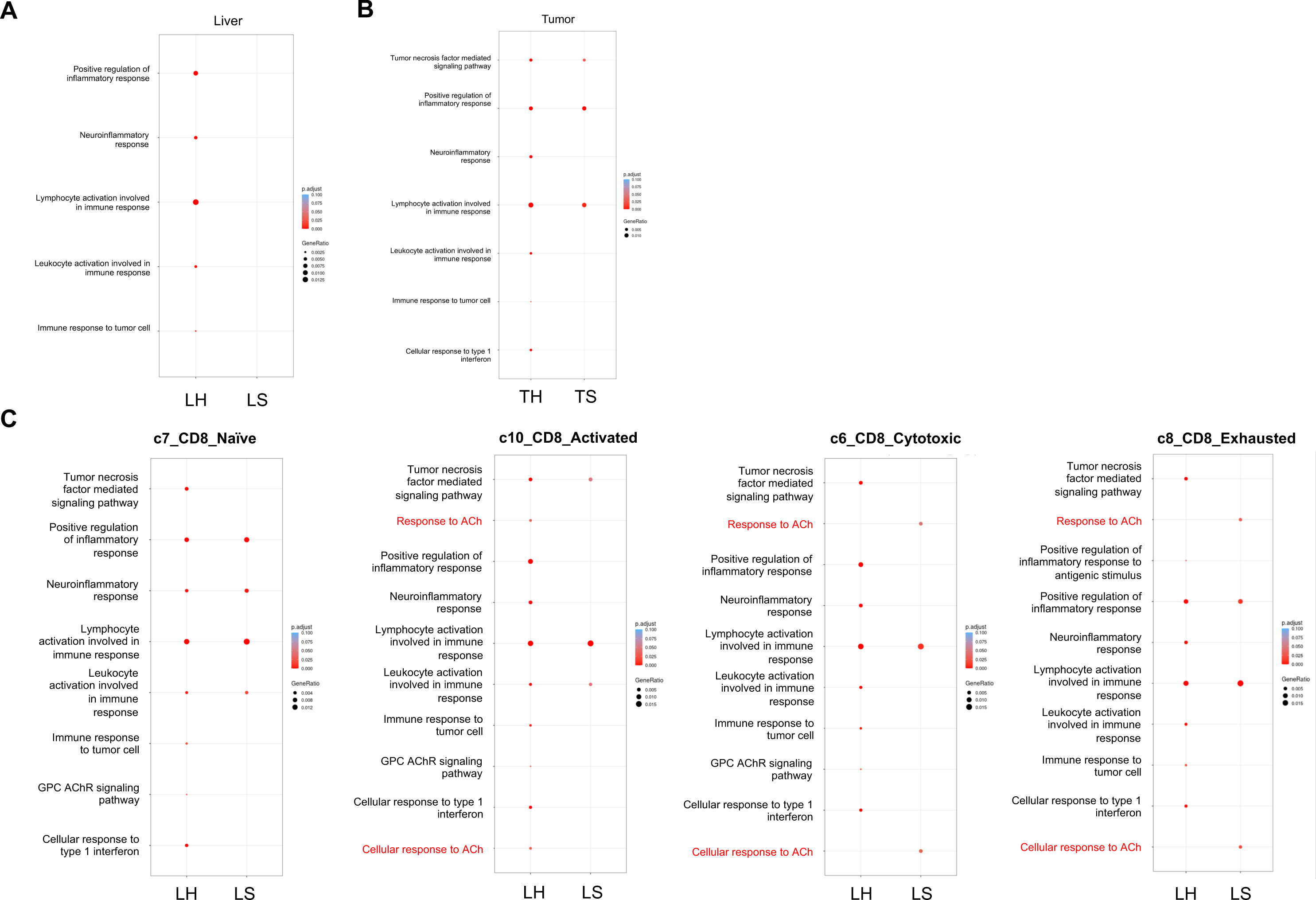
scRNA-Seq identifies distinct ACh regulation in CD8+ T cell clusters. (**A**) 12 pathways were selected as a neuroimmune panel (see **Supplemental File 4**). Pathway enrichment analyses for neuroimmune panel in hepatic vagotomized and sham vagotomized livers, LH and LS, respectively (*left*). (**B**) (*Right*) pathway enrichment analyses for neuroimmune pathways in hepatic vagotomized and sham vagotomized tumors, TH and TS, respectively. Only statistically significant pathways appear in dot plots, color indicates *Padj* and size reflects GeneRatio. Data for all immune (sorted Live+CD45+) cells. (**C**) (*Left-right*) neuroimmune pathway enrichment analyses across CD8+ T cell clusters, see **Fig 3A**: c7_CD8_Naïve, c10_CD8_Activated, c6_CD8_Cytotoxic, and c8_CD8_Exhausted. For scRNA-Seq data, *n* = 3 / group of liver and matched pooled tumor samples.

**Figure S9:**
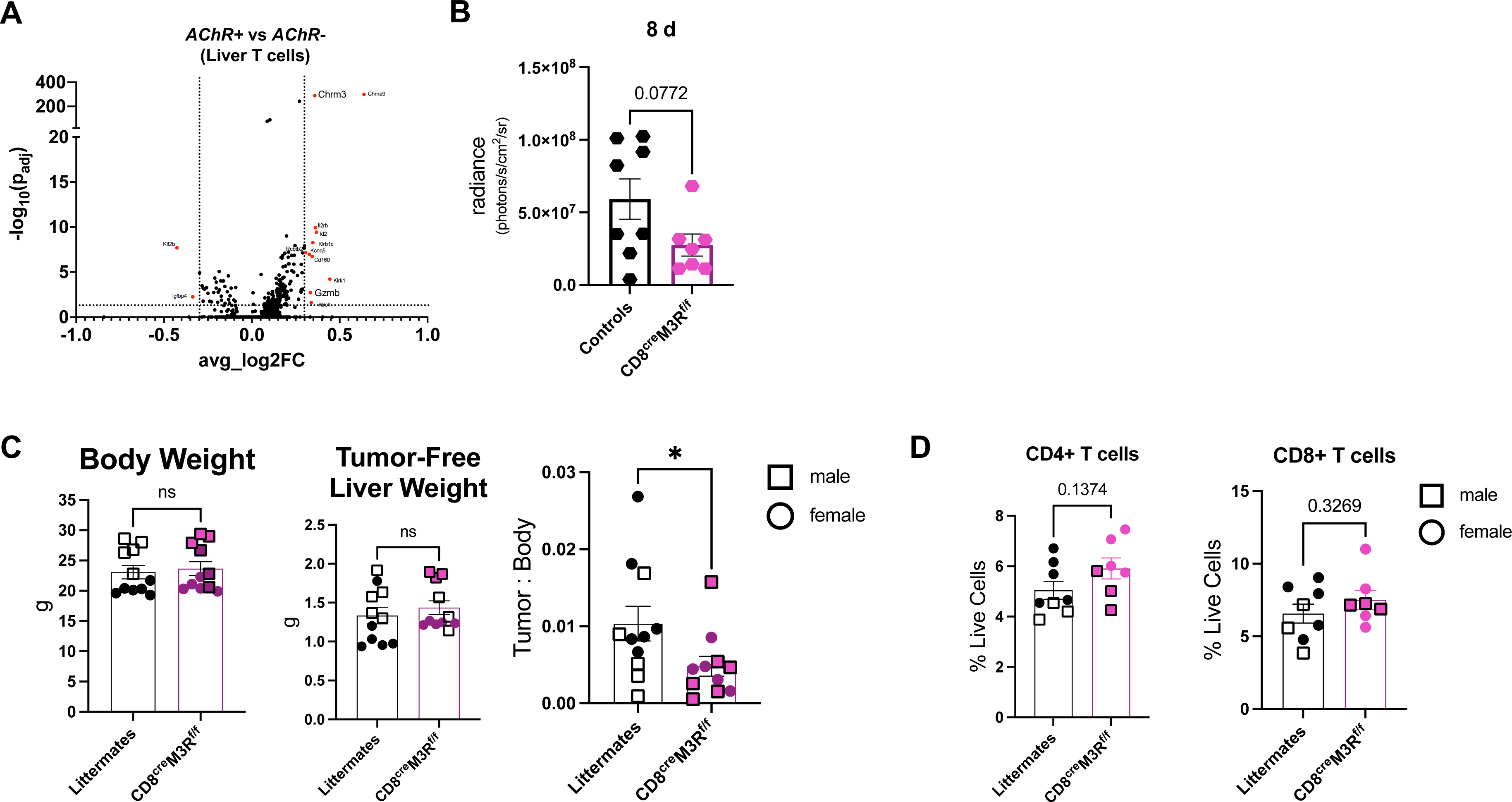
AChR signaling alters hepatic tumor growth and influences CD8+ T cell cytotoxicity. (**A**) Differentially expressed gene (DEG) scRNA-Seq analysis; volcano plot of *AChR*+ vs *AChR-* hepatic T cells in RIL175 model (*n* = 3 / group), significance assessed by 2-way ANOVA. Vertical dotted line reflects 1.5X fold change based on average log2 fold change (log2FC), horizontal dotted line reflects 0.05 *Padj* cutoff. (**B**) Representative in vivo imaging (IVIS® Spectrum) from luciferase-expressing RIL175 tumors 8 d following intrahepatic injection (*n* = 8 CD8^cre^M3R^f/f^ and 7 littermate controls). (**C**) (*Left-right*) total body weight, non-tumorous liver weight, and RIL175 tumor:body ratio in CD8^cre^M3R^f/f^ and littermate controls 14 d after intrahepatic injection, *n* = 11 / group. Data from two independent experiments including cohort represented in **B**. (**D**) Frequency (% live cells) of CD4+ and CD8+ T cells in CD8^cre^M3R^f/f^ and littermate controls, from a subset presented in **C**. Circles reflect female mice and squares indicate male mice. Statistical significance in **B**-**D** determined by unpaired Student’s *t* test. Bar graphs represent mean ± SEM: 0.05<*P*<0.1; \**P* < 0.05; 1, ns = not significant or reported value.

**Figure S10.**
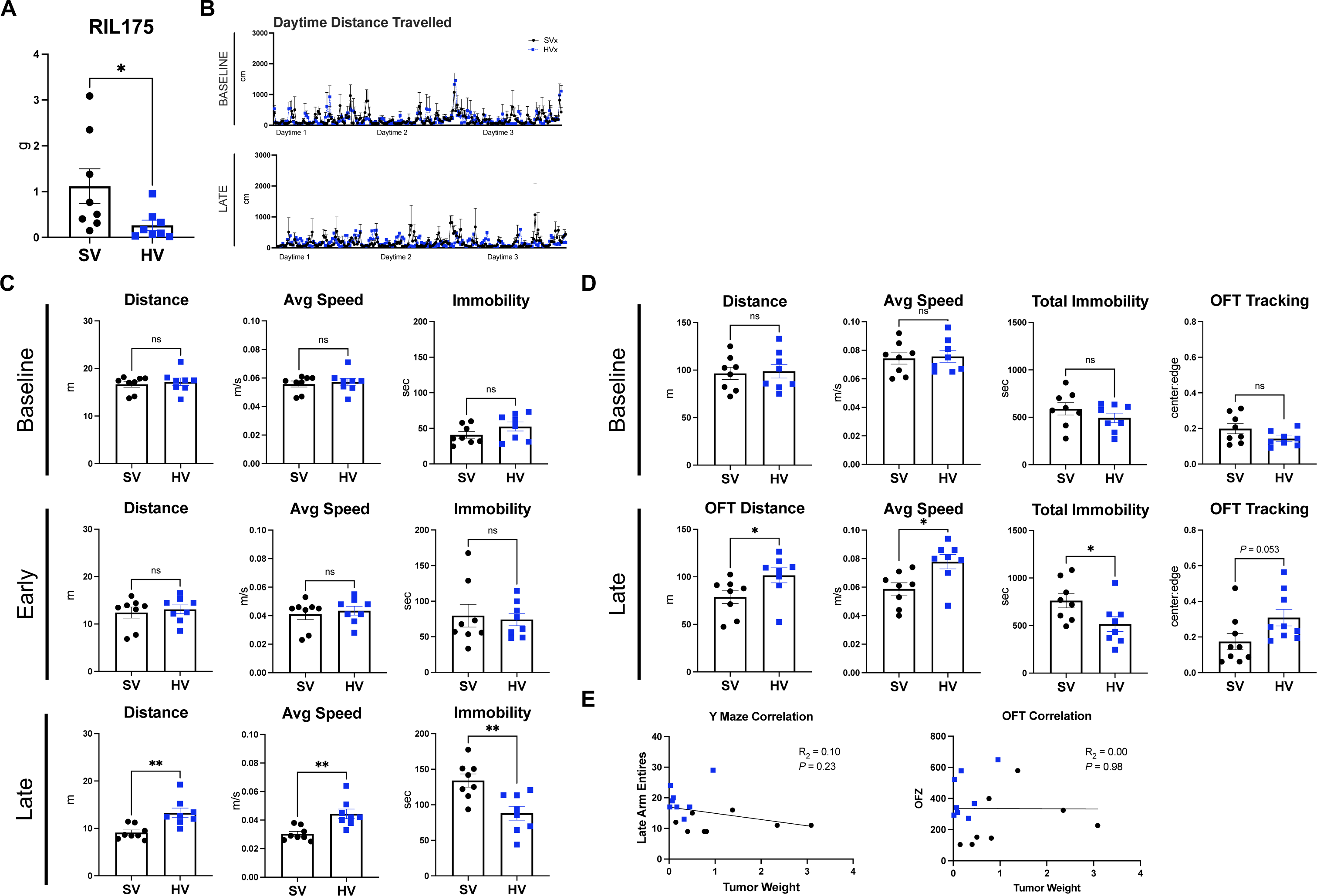
Impaired vagal signaling shapes behavior. (**A**) RIL175 tumor weights in behavioral model 26 d following intrahepatic injection. (**B**) Phenotyper distance (total distance moved from center-point) during daytime tracking at Baseline and Late timepoints (*n* = 3 / group, 10-min intervals recorded), see Fig. 4A. (**C**) Activity assessments for Baseline (post-vagotomy), Early (post-tumor), and Late (post-tumor) timepoints during the Y-maze including total distance traveled, average speed, and total immobility time. (**D**) Activity assessments for Baseline (post-vagotomy) and Late (post-tumor) timepoints during the OFT, including total distance traveled, average speed, and total immobility time, and ratio of time within the OFZ and periphery (center:edge). (**E**) Linear correlation of endpoint tumor weights and late activities: Y maze arm entries (*left*) and OFZ time (*right*). All mice within the panel from the same behavioral study, *n* = 8 / group. Statistical significance determined by linear regression (**E**) or unpaired Student’s *t* test, OFT = open field test, OFZ = open field zone. Bar graphs represent mean ± SEM: reported value 0.05<*P*<0.1, \**P* < 0.05; \*\**P* < 0.01; ns = not significant or reported value.

**Figure S11.**
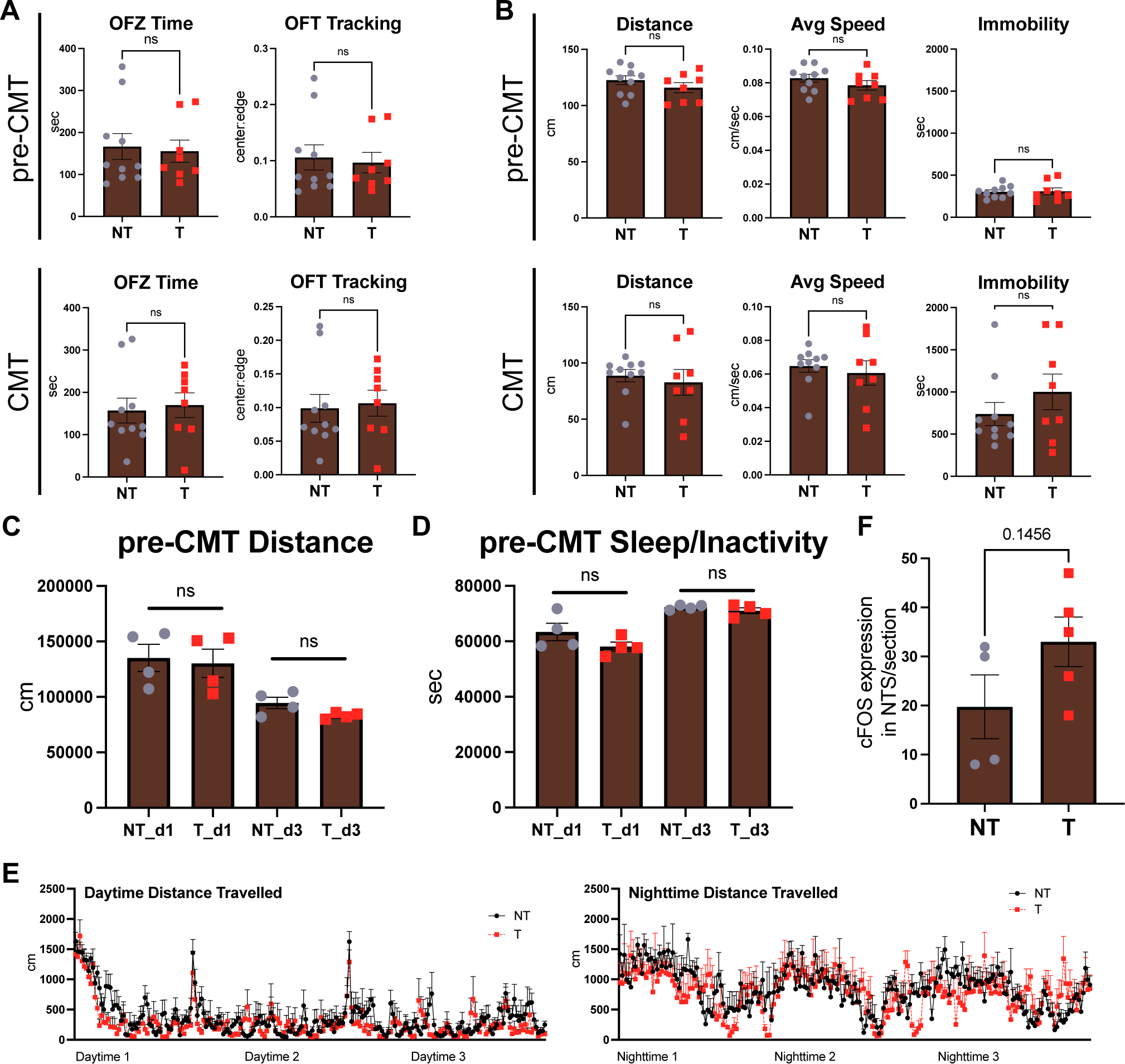
Tumor-bearing CMT alters locomotion. (**A**) OFT results from T (tumor) and NT (non-tumor) cecal microbiota transplant (CMT) study, see Fig. 4G. Total OFZ time and ratio of time spent in the OFZ to peripheral zone (center:edge) prior to CMT (*above*) and following CMT (*below*). (**B**) OFT activity pre-CMT (*above*) and post-CMT (*below*) including total distance traveled, average speed, and total immobility time, *n* = 10 NT, 8 T mice. (**C**) Phenotyper distance (total distance moved from center-point) and (**D**) immobility (sleep/inactivity periods) in a subset of mice prior to CMT (*n* = 4 / group). Due to a software malfunction on the second day, only data from the 1^st^ and 3^rd^ session are reported. Data averaged across 12 h. PhenoTyper 3000 cage availability determined *n* size in CMT experiment, remaining mice were singly caged during phenotyping analyses (**E**) Daytime and nighttime phenotyper distance (total distance moved from center-point) from post-CMT testing (*n* = 7 NT, 5 T). (**F**) cFOS counts per NTS region in CON and RIL175 mice, staining 14 d after tumor initiation (*n* = 4 NT, 5 T). Except for **F**, mice in the S9 panel from the same CMT study; OFT = open field test, OFZ = open field zone. Graphs represent mean ± SEM. Statistical significance determined by unpaired Student’s *t* test: ns = not significant.

**Figure S12.**
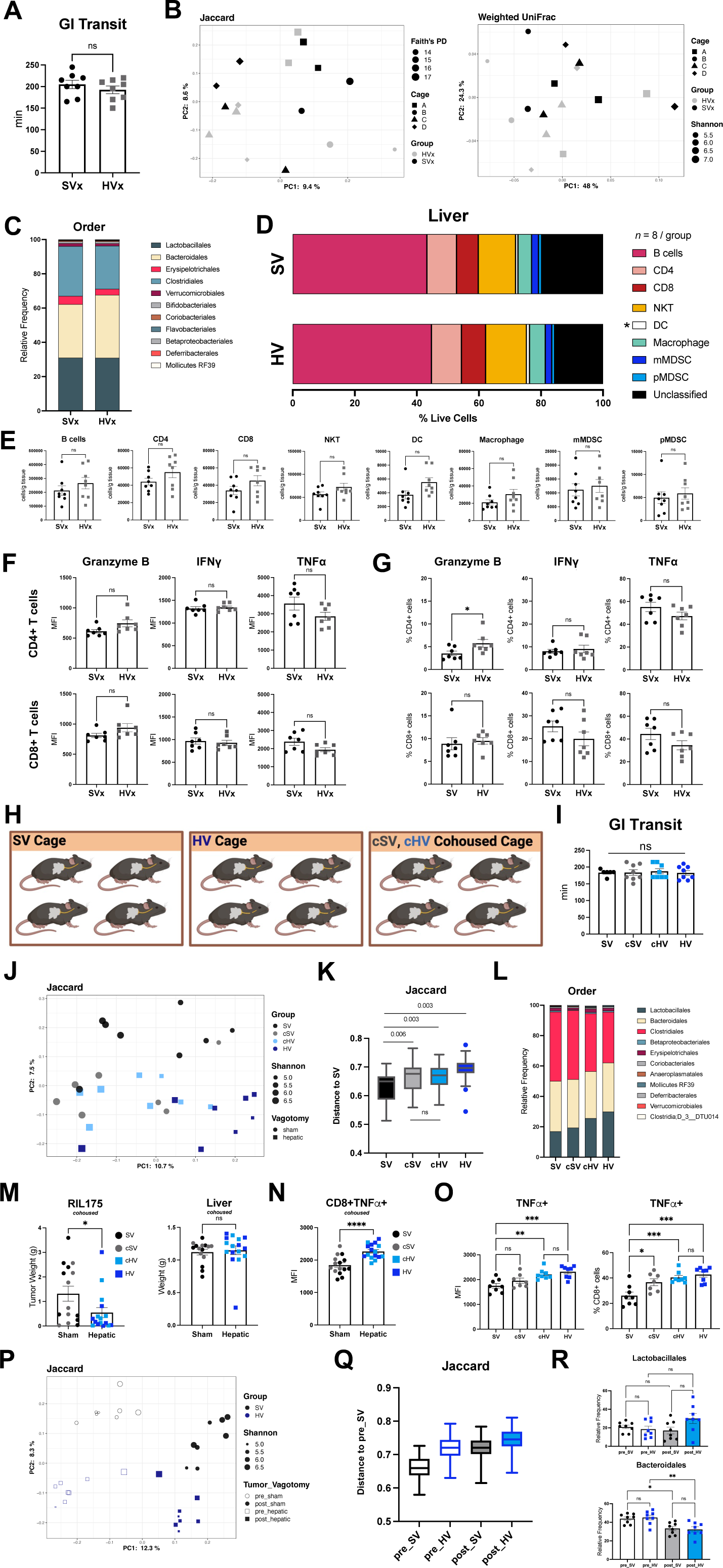
Anti-tumor immunity not linked to vagotomized microbiome. (**A**) GI (gastrointestinal) transit time as measured via carmine red test. (**B**) Beta diversity (Jaccard, Weighted UniFrac) and alpha diversity (Faith’s PD, Shannon entropy) from fecal microbiota samples of non-tumor bearing cohoused mice that underwent a hepatic vagotomy (HVx) or sham surgery (SVx). Following vagotomy, mice were randomized and cohoused for three weeks. Jaccard (*left*) and Weighted UniFrac (*right*) plots determined following 16S rRNA-Seq of fecal samples. (**C**) Bacterial composition (Order) in SVx and HVx mice, *n* = 8 / group. Mice in **A-C** from the same cohort, see also **Supplemental File 5**. (**D**) Immune profiling in SVx and HVx mice assessed by flow cytometry, frequency of Live+ hepatic cells presented. (**E**) Cell counts of Live+ immune cells normalized to tissue weights from hepatic SVx and HVx samples (*n* = 14 / group). (**F**) Granzyme B, IFNγ+ and TNFα+ staining in *ex vivo* lymphocytes following 4 h PMA/ionomycin stimulation, MFI and (**G**) parental frequency reported for CD4+ and CD8+ T cells, n = 7 / group. (**H**) Caging strategy for cohousing tumor experiment. Post vagotomy, HVx and SVx mice were randomized into non-cohoused (SV or HV) or cohoused cages (cSV and sHV mice) prior to RIL175 tumor initiation. Tumor weights and final fecal microbiota samples were assessed at 21 d. For cohousing experiment reported in **I**-**R**, *n* = 8 SV, 7cSV, 8 cHV, and 8 HV. (**I**) GI transit time prior to cohousing termination in tumor-bearing SV, cSV, sHV, and HV mice assessed via carmine red test. (**J**) Jaccard beta diversity plot of SV, cSV, cHV, and HV fecal samples with Shannon entropy (alpha diversity) reported. (**K**) Jaccard distances from **K**, pairwise PERMANOVA with q-value reported. (**L**) Bacterial composition determined via 16S rRNA-Seq from fecal samples, Order reported. (**M**) (*Left*) tumor weight and (*right*) non-tumor liver weights at 21 d in cohousing experiment, color indicative of group and shape identifies vagotomy status: circle = sham procedure, square = hepatic vagotomy. (**N**) *Ex vivo* intracellular cytokines staining in hepatic tissue following stimulation reported in (**F**). MFI of CD8+TNFα+ mice. (**O**) Both the MFI (*left*) and frequency (*right*) of CD8+TNFα+ shown in **N** separated by cohoused and non-cohoused groups. (**P**) Beta (Jaccard) and alpha (Shannon entropy) plot from 16S rRNA-Seq assessment of SV and HV fecal samples prior to liver cancer initiation (Tumor_Vagotomy = pre_SV, pre_HV) and at experimental termination (Tumor_Vagotomy = post_SV, post_HV). SV and HV beta diversity composition reported in **J**. (**Q)** Jaccard distances from **P** (set to pre_SV), pairwise PERMANOVA performed and all q-values <0.001. (**R**) Bacterial composition determined via 16S rRNA-Seq from fecal samples, top two taxa (Order) reported in pre- and post-tumor timepoints. Bar graphs (**A**, **E**-**G**, **I**, **M**-**O**, and **R**) represent mean ± SEM or boxplot (min-max whiskers): \**P* < 0.05; \*\**P* < 0.01; \*\*\**P* < 0.005; \*\*\*\**P* < 0.001, ns = not significant.

**Figure S13.**
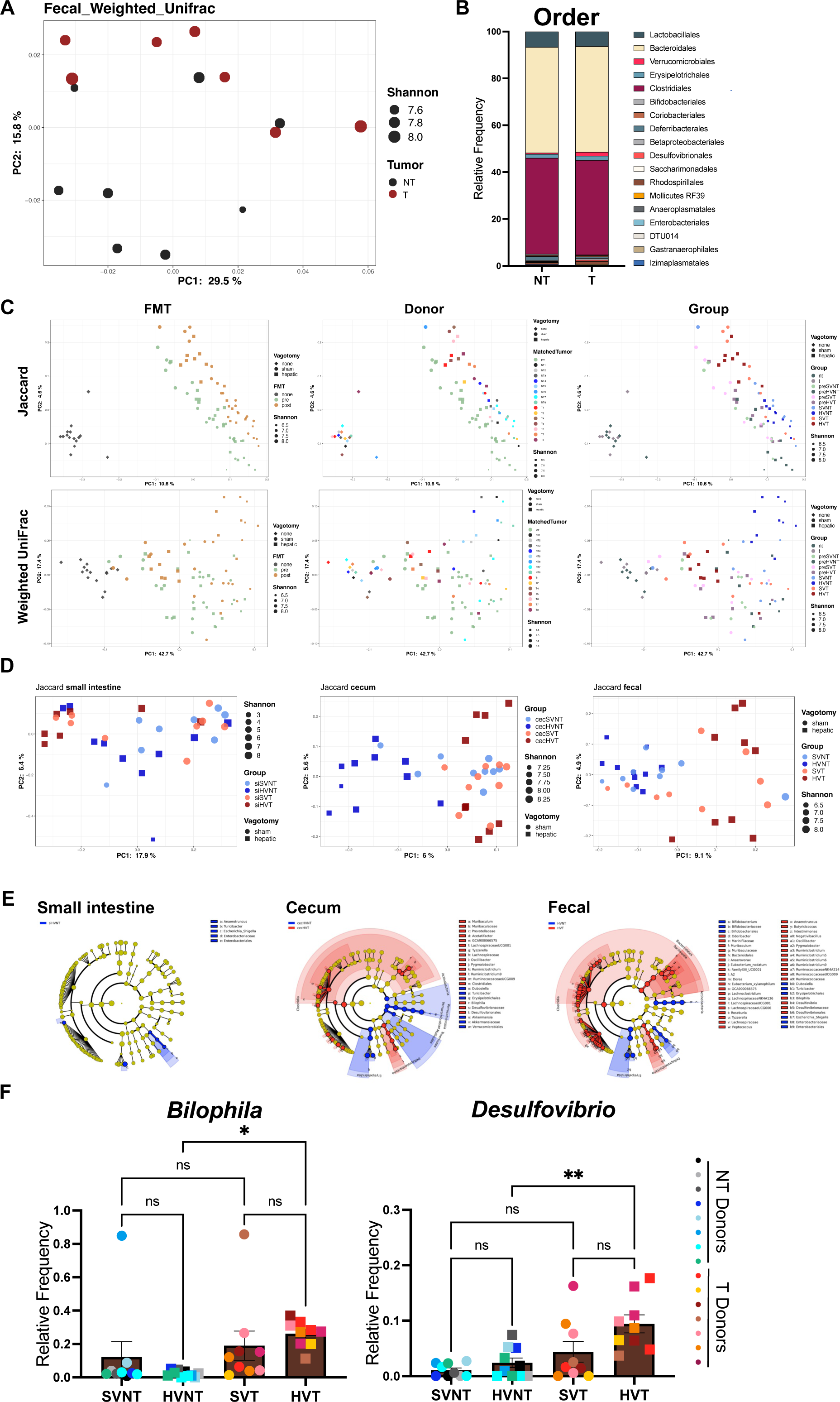
Tumor status rather than vagotomy drives CMT findings. (**A**) Fecal microbiota analyses from 16S rRNA-Seq of frozen cecal content from tumor (T) and non-tumor (NT) donors, alpha diversity (Shannon entropy) and beta diversity reported (Weighted UniFrac) in cecal microbiota transplant (CMT) study. (**B**) Bacterial composition (Order) from NT and T donor repository. (**C**) Timeline plots of alpha (Shannon entropy) and beta diversity: (*above*) Jaccard, (*below*) Weighted UniFrac assessing pre-CMT microbiota and post-CMT (final) fecal collection. From *left*-*right* color indicates timepoint collection (none = donor, pre = pre-CMT fecal sample, post = final fecal collection), donor status (unique donor CMT), and group (*i.e.,* SVNT, HVNT, SVT, and HVT). For CMT analyses, Donors: *n* = 8 NT, 7 T; Recipients: *n* = 9 SVNT, 10 HVNT, 9 SVT, and 9 HVT. (**D**) Jaccard beta diversity with Shannon entropy (alpha diversity) from small intestine, cecal, and fecal samples, Weighted UniFrac presented in Fig. 5C. (**E**) LefSE analyses (*84*) of HVNT and HVT fecal microbiota samples with differentially expressed taxa visualized via cladogram from (*left*-*right*) small intestine, cecum, and fecal samples. For LefSE analyses across the GI tract: α = 0.01 Kruskal-Wallis, α = 0.01 Wilcoxon test, LDA > 2 (less stringent than Fig. 5F). No enriched bacterial taxa were reported in SVT and HVT fecal samples using these LefSe settings. (**F**) Relative frequency of bacterial species (*left*) *Bilophila* and (*right*) *Desulfovibrio* from fecal samples. Statistical significance in **F** determined by one-way ANOVA with *posthoc* Tukey testing. Bar graphs in **F** represent mean ± SEM: \**P* < 0.05; \*\**P* < 0.01; ns = not significant.

**Figure S14:**
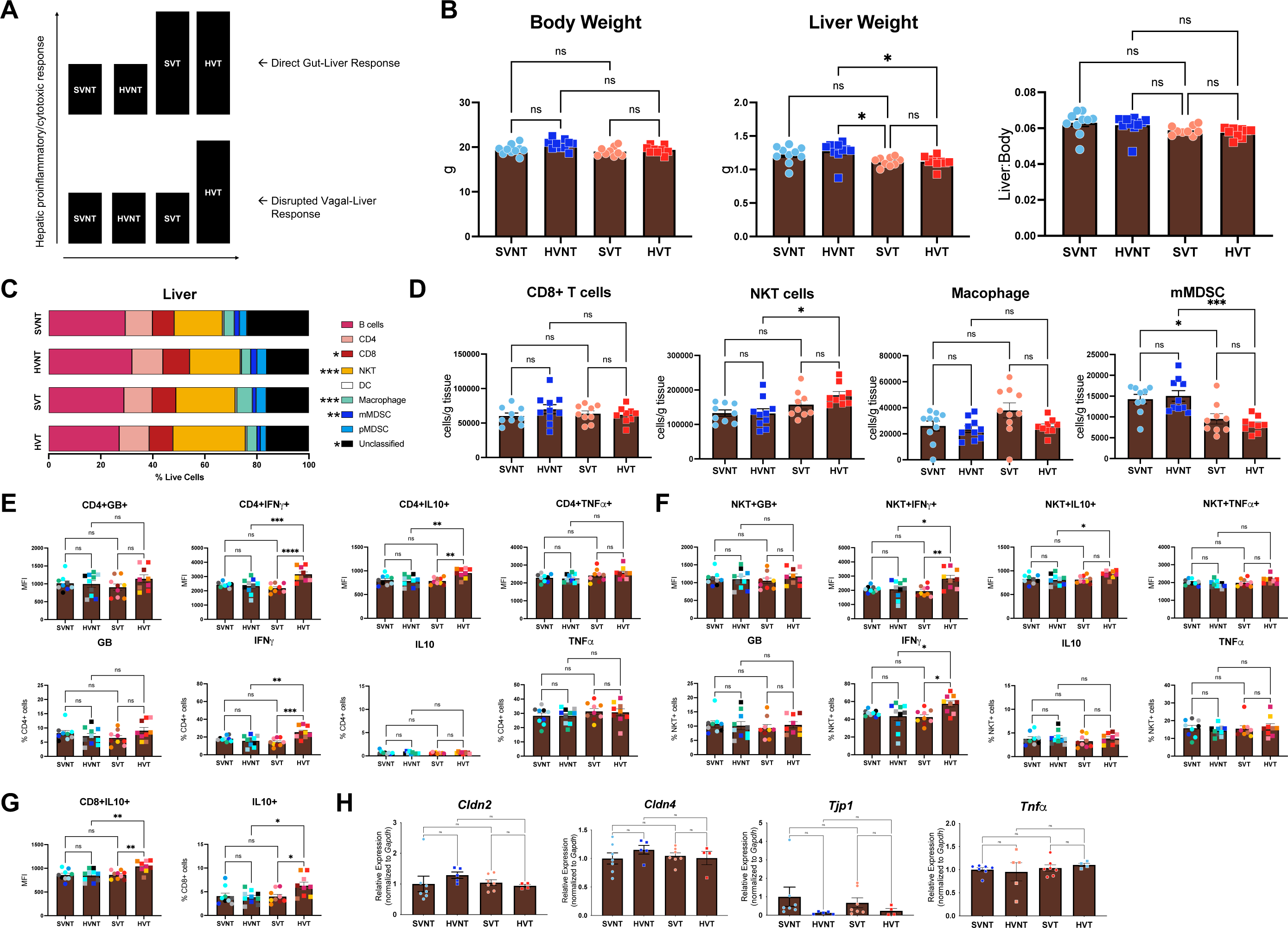
Vagal-liver axis alters liver immune composition and function. (**A**) Model of putative gut ➔ liver (*above*) and vagus ➔ liver (*below*) hepatic responses in CMT study. X-axis reports recipient group and Y-axis reflects hepatic inflammatory or CD8+ T cell effector function. While T microbiota provokes hepatic response via gut-liver interactions (top), we propose that SVT mice maintain liver immunosuppressive-like features via vagal integrity, *i.e.*, vagus ➔ liver pathway (bottom). (**B**) Gross body measurements in CMT study, (*left*-*right*) total body weight, liver weight, and liver:body ratio. (**C**) Immunoprofiling assessed via flow cytometry, reporting % Live+ hepatic cells. (**D**) Cell counts of select flow cytometry data in **C** normalized to tissue weight, (*left*-*right*) CD8+ T cells (CD3+CD8+), NKT cells (CD3+CD1d tetramer+), macrophage ([non-mMDSC] CD3-CD11b+F4/80+), and mMDSC populations (CD3-CD11b+F4/80+Ly6C+Ly6G-). (**E**) *Ex vivo* intracellular cytokine staining after PMA/ionomycin stimulation as reported in Expanded Methodology. (*Above*) MFI and (*below*) parental frequency of cytokine markers in CD4+ cells and (**F**) NKT+ cells. (**G**) Parental frequency and MFI of CD8+ markers, IL10+ staining reported. (**H**) RT-qPCR data (left-right) *Cldn2*, *Cldn4*, *Tjp1*, and *Tnf*α expression from proximal colon (10 cm). Data normalized to *Gapdh*, reporting fold change from SVNT relative expression. Statistical significance determined by one-way ANOVA with *posthoc* Tukey’s multiple comparisons; *Cldn* = claudin, *Tjp* = tight junction protein. Bar graphs represent mean ± SEM: \**P* < 0.05; \*\**P* < 0.01; \*\*\**P* < 0.05; \*\*\*\**P* < 0.001; ns = not significant.

## Supplemental File Legends

**Supplemental File 1 Tumor adjacent tissue exhibits altered immune and neurological pathways**

Pathway enrichment analyses outputs for all cancer types

**Supplemental File 2 CODEX identification markers**

18-color antibody panel for identification of nearest cellular neighbors

**Supplemental File 3 scRNA-Seq differentially expressed gene list**

DEG from all liver and tumor scRNA-Seq immune cells ranked by average log fold change value

**Supplemental File 4 Neuroimmune gene list**

KEGG GO values for the Neuroimmune Panel

**Supplemental File 5 Microbiome Analyses**

Further beta and alpha diversity statistics and analyses for cohousing and CMT experiments.

## Notes

### Competing Interest Statement

The authors have declared no competing interest.

