## Supplemental File 5 Microbiome Analyses for "The Gut Microbiome Controls Liver Tumors via the Vagus Nerve"

Unweighted UniFrac distances for cohoused non-tumor bearing mice (*n* = 7 SVx, 7 HVx). Distances by group. Distance to SVx (left) and HVx (right). PERMANOVA: F = 0.9129, *P* = 0.59.

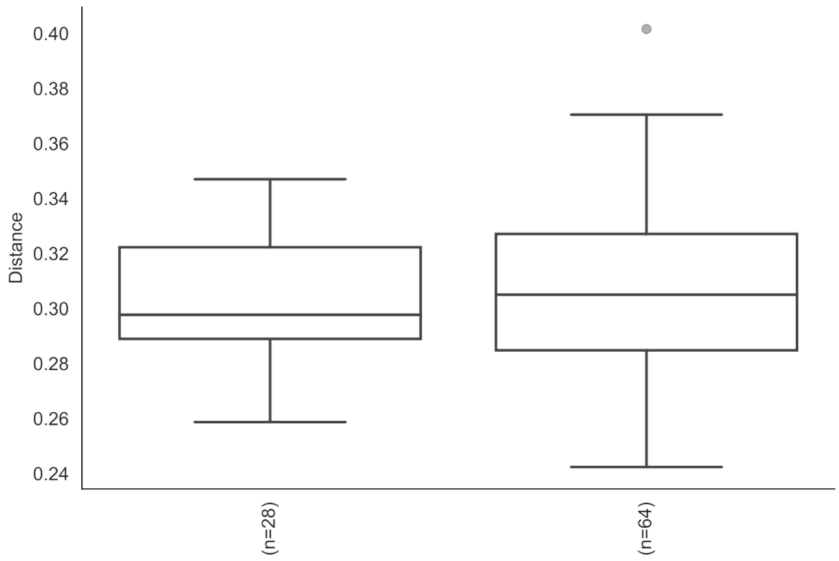


Unweighted UniFrac distances for cohoused non-tumor bearing mice (*n* = 7 SVx, 7 HVx). Distances by cage. Distance to Cage “A”. PERMANOVA: F = 1.193721, *P* = 0.001.

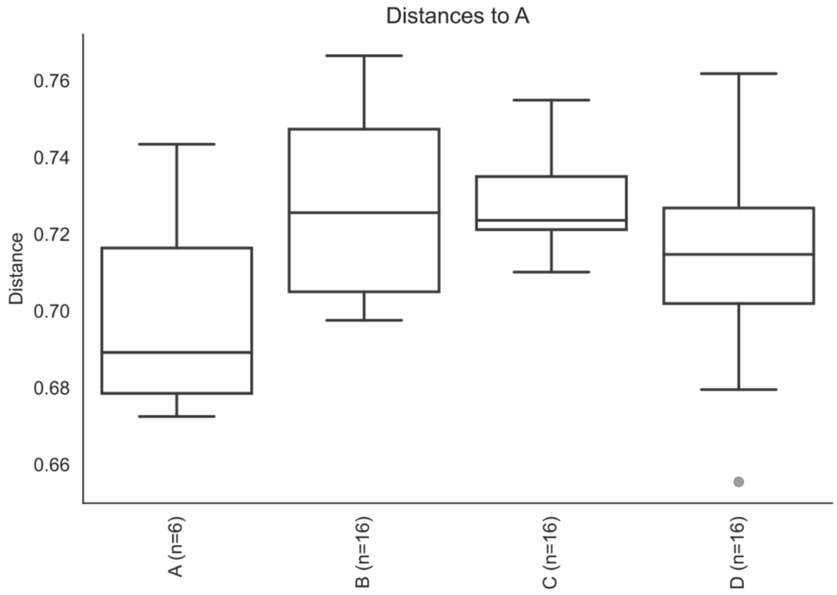

Jaccard distances for cohoused experiment, final fecal collection (n = 8 SV, 7 cSV, 8 cHV, and 8 HV). PERMANOVA: F = 1.769972, *P* = 0.001.

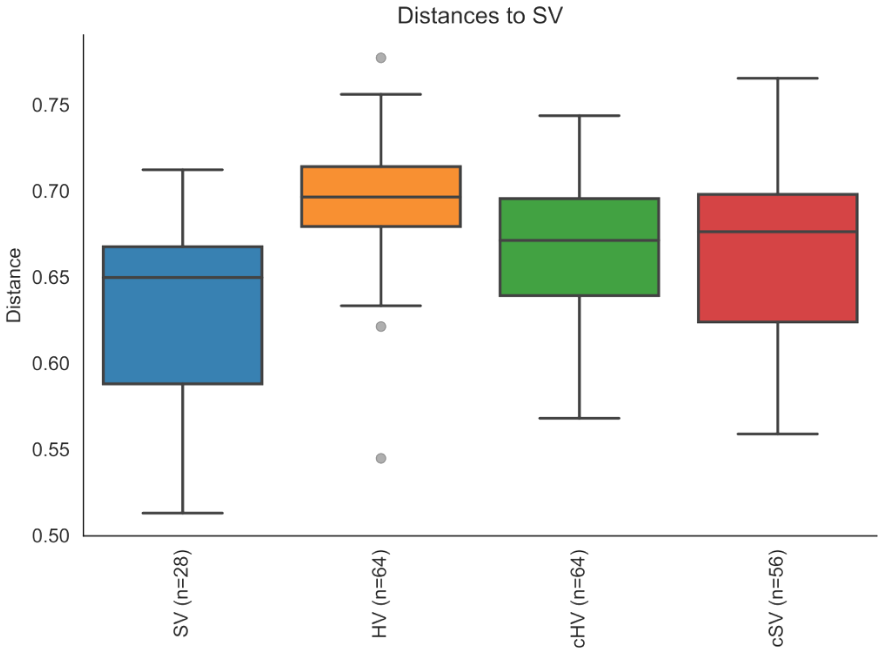


Jaccard distances for cohoused experiment, “pre-post”, paired SV and HV samples “pre” (vagotomy, pre-tumor) and “post” (vagotomy, tumor) (n = 8 SV and 8 HV). PERMANOVA: F = 2.9866, *P* = 0.001. See sFig10 for distances against pre_SV.

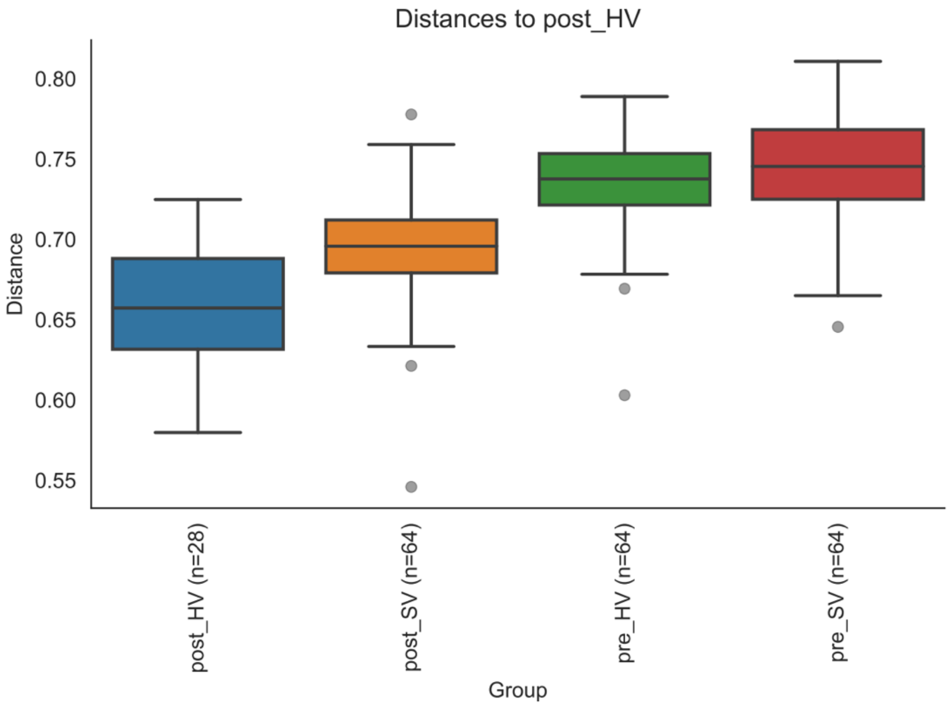

Pairwise distance analyses (Jaccard) distances for cohoused experiment.

### Multiple group tests

| | H | P value |
|---|---|---|
| **Kruskal Wallis test** | 1.588235 | 0.207578 |

### Pairwise group comparison tests

| | | Mann-Whitney U | P-value | FDR P-value |
|---|---|---|---|---|
| **Group A** | **Group B** | | | |
| **HV** | **SV** | 44.0 | 0.234499 | 0.234499 |

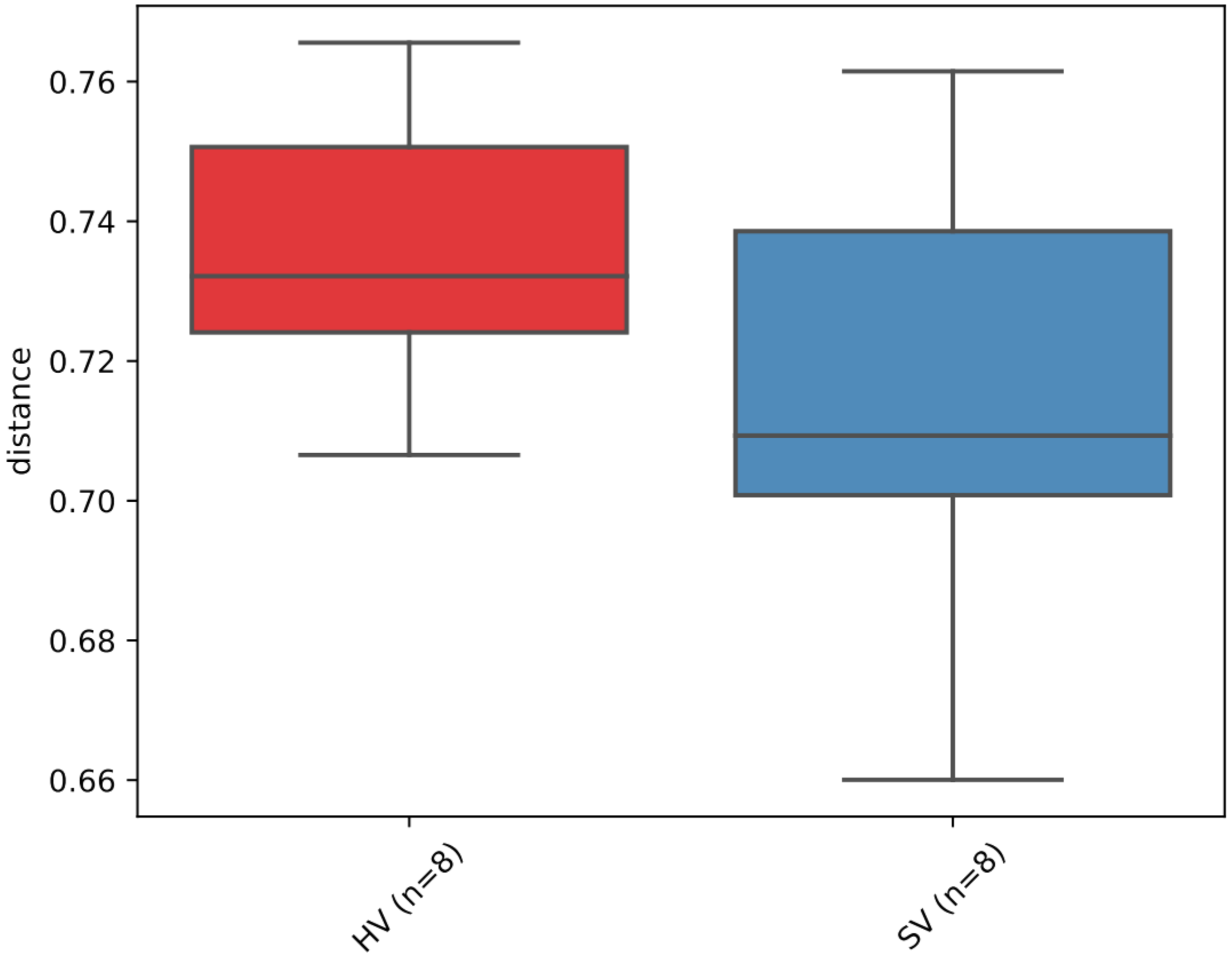

Jaccard distances for CMT experiment (n = 9 SVNT, 10 HVNT, 9 SVT, 9 HVT). Ileum samples. Distances based on tumor status (*top*). PERMANOVA: F = 1.55458, *P* = 0.03 and vagotomy status (*bottom*). PERMANOVA: F = 1.74171, *P* = 0.026.

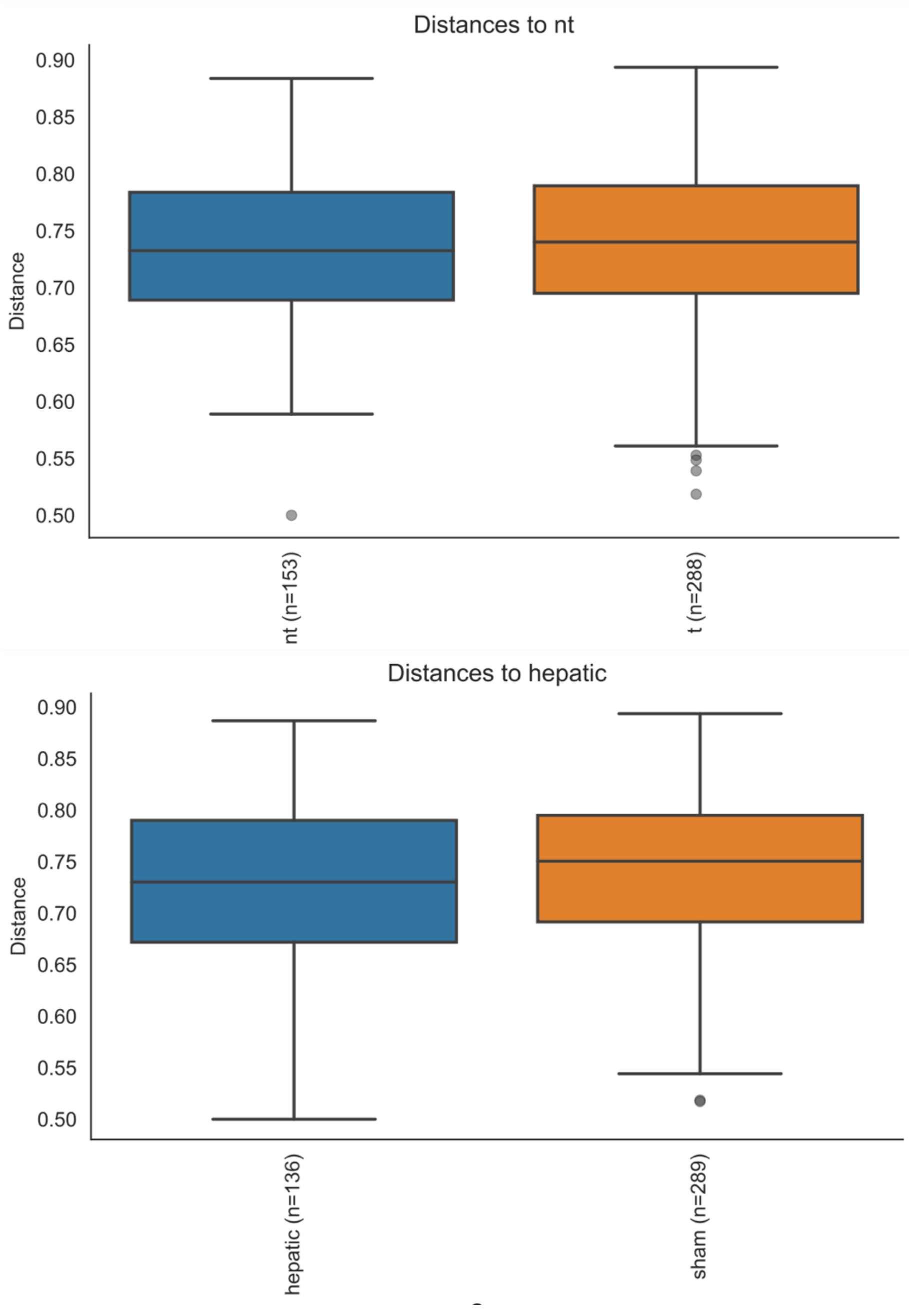

CMT Cecal samples. Distances based on tumor status (*top*). PERMANOVA: F =1.581064, *P* = 0.001 and vagotomy status (*bottom*). PERMANOVA: F = 1.423151, *P* = 0.002.

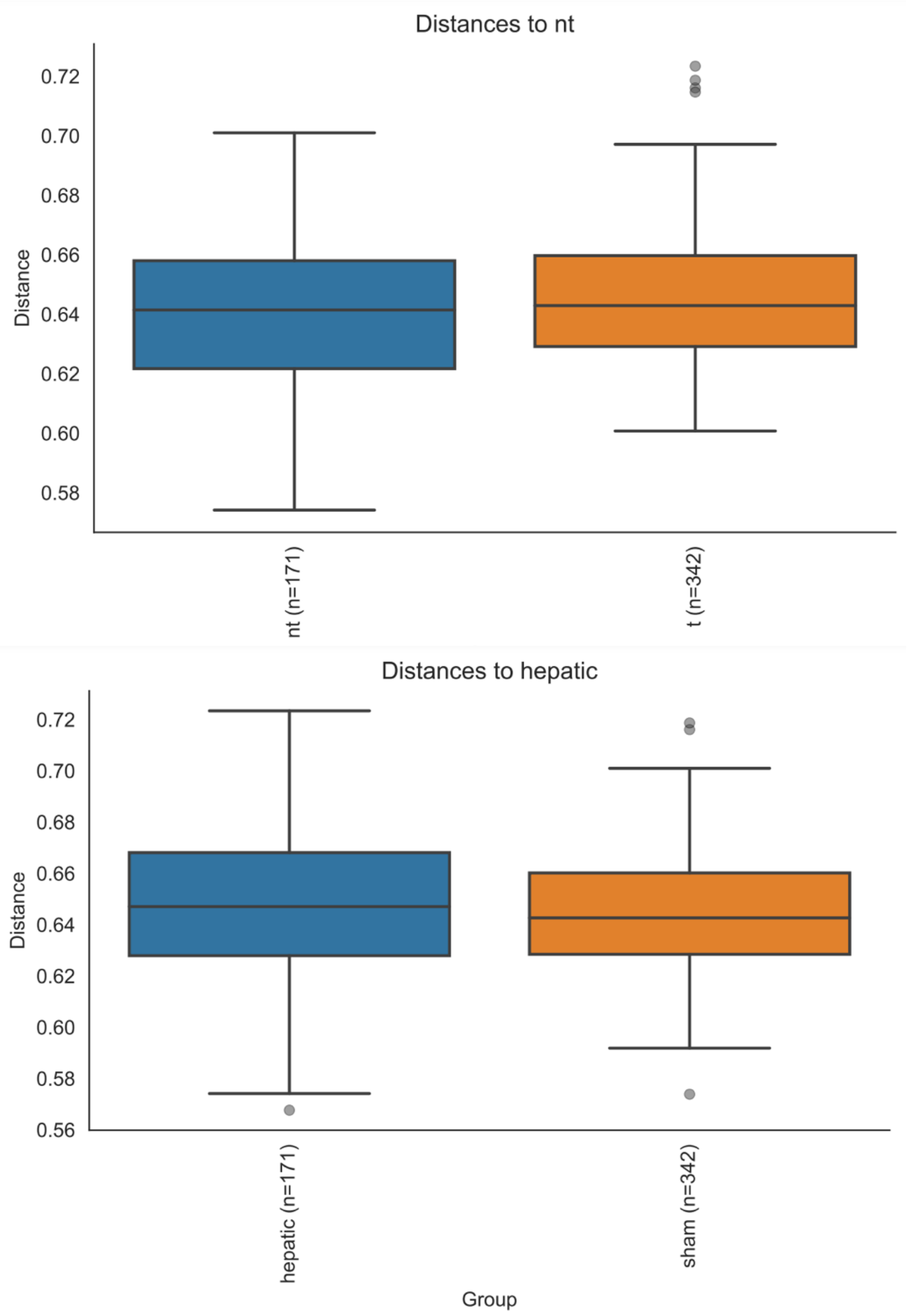

CMT Fecal samples. Distances based on tumor status (*top*). PERMANOVA: F = 2.03457, *P* = 0.001 and vagotomy status (*bottom*). PERMANOVA: F = 1.076221, *P* = 0.185.

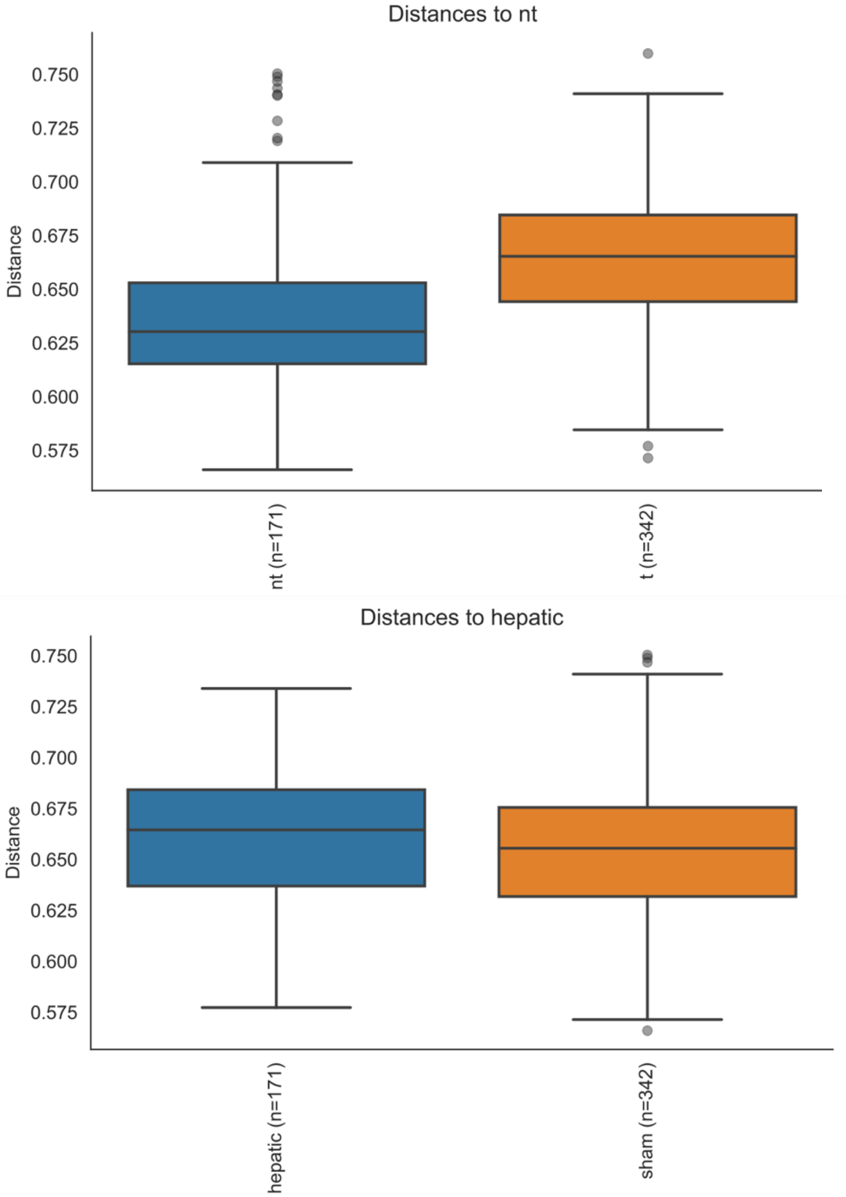
